# A genomic duplication spanning multiple P450s contributes to insecticide resistance in the dengue mosquito *Aedes aegypti*

**DOI:** 10.1101/2024.04.03.587871

**Authors:** Tiphaine Bacot, Chloé Haberkorn, Joseph Guilliet, Julien Cattel, Mary Kefi, Louis Nadalin, Jonathan Filee, Frederic Boyer, Thierry Gaude, Frederic Laporte, Jordan Tutagata, John Vontas, Isabelle Dusfour, Jean-Marc Bonneville, Jean-Philippe David

**Affiliations:** Laboratoire d’Ecologie Alpine (LECA, UMR 5553), Université Grenoble-Alpes (UGA), Université Savoie Mont-Blanc (USMB), CNRS, 38041 Grenoble, France; Université Paris-Saclay, CNRS, IRD, UMR Évolution, Génomes, Comportement et Écologie, 91198 Gif-sur-Yvette, France; Institute of Molecular Biology and Biotechnology, Foundation for Research and Technology Hellas, Heraklion, Greece; Pesticide Science Laboratory, Department of Crop Science, Agricultural University of Athens, Athens, Greece; Vectopôle Amazonien Emile Abonnenc, Institut Pasteur de la Guyane, Cayenne, France; Global Health Department, Institut Pasteur, Paris, France

**Keywords:** Mosquito, Insecticide resistance, P450, Gene duplication, Kdr mutation, *Aedes aegypti*

## Abstract

Resistance of mosquitoes to insecticides is one example of rapid adaptation to anthropogenic selection pressures having a strong impact on human health and activities. Target-site modification and increased insecticide detoxification are the two main mechanisms underlying insecticide resistance in mosquitoes. While target-sites mutations are well characterised and often used to track resistance in the field, the genomic events associated with insecticide detoxification remain partially characterised. Recent studies evidenced the key role of gene duplications in the over-expression of detoxification enzymes and their potential use to track metabolic resistance alleles in the field. However, such genomic events remain difficult to characterise due to their complex genomic architecture and their co-occurrence with other resistance alleles. In this concern, the present work investigated the role of a large genomic duplication affecting a cluster of detoxification enzymes in conferring resistance to the pyrethroid insecticide deltamethrin in the mosquito *Aedes aegypti*.

Two isofemale lines originating from French Guiana and being deprived from major target-site mutations showed distinct insecticide resistance levels. Combining RNA-seq and whole genome pool-seq identified a 220 Kb genomic duplication enhancing the expression of multiple contiguous cytochrome P450s in the resistant line. The genomic architecture of the duplicated loci was elucidated through long read sequencing, evidencing its transposon-mediated evolutionary origin. The involvement of this P450 duplication in deltamethrin survival was supported by a significant phenotypic response to the P450 inhibitor piperonyl butoxide together with genotype-phenotype association and RNA interference. Experimental evolution suggested that this P450 duplication is associated with a significant fitness cost, potentially affecting its adaptive value in presence of other resistance alleles.

Overall, this study supports the importance of genomic duplications affecting detoxification enzymes in the rapid adaptation of mosquitoes to insecticides. Deciphering their genomic architecture provides new insights into the evolutionary processes underlying such rapid adaptation. Such findings provide new tools for the surveillance and management of resistance in the field.

## Introduction

Natural populations experience a wide range of selective pressures, leading to the accumulation of locally adaptive features and the expression of complex phenotypes (Orr, 2005). Among them, those driven by human activities promote novel and strong selective pressures. Understanding the genetic mechanisms allowing populations to quickly respond to rapid environmental changes has become a major goal (Hendry et al., 2008, 2017; Palumbi, 2001). Resistance of insects to insecticides is a key example of rapid evolution under strong anthropogenic selective pressures. This adaptation has occurred quickly and independently in a large number of taxa with frequent parallel trajectories (Ffrench-Constant et al., 2004; Kamdem et al., 2017; Liu, 2015). As a consequence of local and temporal variations of the selection pressures, the genetic modifications affecting resistant populations are often complex and combine several resistance mechanisms that are potentially additive, each of them bringing variable fitness costs. In addition, the very strong selection pressure exerted by insecticides is prone to produce bottleneck effects. Such complexity makes the identification of resistance alleles and the assessment of their respective contribution challenging (Ffrench-Constant et al., 2004; Li et al., 2007).

Besides the understanding of the genetics of rapid adaptation and the origins of complex traits, deciphering the molecular bases of insecticide resistance is also essential for improving pest management strategies (Hawkins et al., 2018). Among taxa of major economic and medical importance, mosquitoes represent a major threat for public health worldwide because of their ability to transmit human viruses and pathogens (Lounibos, 2002). Among them, *Aedes aegypti* is of particular importance because of its wide distribution and its capacity to transmit Yellow fever, Dengue, Zika and Chikungunya viruses (Brown et al., 2014). These arboviruses are now re-emerging worldwide following the expansion of mosquitoes’ distribution area as a consequence of global warming, global transportation network and land perturbations (Kraemer et al., 2015). Although efforts are invested in improving vaccine-based prevention strategies (Carvalho & Long, 2021; Garg et al., 2020; Silva et al., 2018), vector control remains the cornerstone of arboviral diseases control. However, decades of insecticide usage have led to the selection and spread of insecticide resistance in this mosquito species, affecting all public health insecticides, including the most-used pyrethroids (Moyes et al., 2017). High pyrethroid resistance has been shown to reduce vector control efficacy (Dusfour et al., 2011; Marcombe, Carron, et al., 2009; Marcombe et al., 2011; Valle et al., 2019).

Though greener vector control strategies are being developed, insecticides will likely remain a key component of integrated vector control in high transmission areas for the next decades (Achee et al., 2019). In this concern, identifying the genetic factors underlying resistance is crucial for tracking resistance alleles in the field and making a better use of the few authorised public health insecticides through resistance management actions (Cattel et al., 2020; Corbel & N’Guessan, 2013; Dusfour et al., 2019).

In mosquitoes, resistance to chemical insecticides is mainly caused by genetic changes decreasing the affinity of the insecticide for its target (target-site resistance), decreasing its penetration (cuticular resistance), or increasing its detoxification through complex biochemical pathways (metabolic resistance) (Li et al., 2007). Pyrethroid insecticides target the neuronal voltage-gated sodium channel (*VGSC* gene), and the selection of *knockdown resistance* (*Kdr*) mutations affecting this protein can lead to resistance. Multiple *Kdr* mutations, often combined as haplotypes, have been identified in *Ae. aegypti* with the following ones playing a major role in pyrethroid resistance in South America: Val410Leu, Val1016Ile and Phe1534Cys (Brengues et al., 2003; Haddi et al., 2017; Hirata et al., 2014; Kasai et al., 2022; Saavedra-Rodriguez et al., 2007; Smith et al., 2016; Yanola et al., 2011). Today, these mutations can be tracked in the field using PCR-based diagnostic assays or mass sequencing approaches, providing essential information for resistance management programmes (Melo Costa et al., 2020).

Conversely, the genetic bases of metabolic resistance are far less understood, though it often accounts for a significant part of the phenotype (David et al., 2013). Indeed, the complexity and redundancy of xenobiotic biodegradation pathways often lead to multigenic adaptations that can differ according to the nature and intensity of the selection pressure together with the demographic and ecological context (Feyereisen et al., 2015; Li et al., 2007). Metabolic resistance to pyrethroids usually results from an increased activity of detoxification enzymes such as cytochrome P450 monooxygenases (P450s or *CYPs* for genes), glutathione S-transferases (GSTs) carboxy/cholinesterases (CCEs) and UDP-glycosyl-transferases (UDPGTs) (David et al., 2013; Hemingway et al., 2004; Smith et al., 2016). At the genetic level, this can result from the selection of enzyme variants showing a higher insecticide metabolism rate, or their overexpression through cis-or trans-regulation. Genomic duplications can also contribute to overexpression, and duplications affecting detoxification genes appear frequently associated with insecticide resistance in mosquitoes (Cattel et al., 2020, 2021; Faucon et al., 2015; Weetman et al., 2018).

In addition to the genetic complexity of metabolic resistance alleles, their frequent co-occurrence with other resistance mechanisms in natural mosquito populations makes them difficult to characterise. This is the case in French Guiana, where the use of various insecticides, including the pyrethroid deltamethrin, for decades has led to the selection of highly resistant *Ae. aegypti* populations combining metabolic resistance alleles and multiple *Kdr* mutations such as Val410Leu, Val1016Ile or Phe1534Cys (Dusfour et al., 2015; Haddi et al., 2017). Although the over-expression of several detoxification enzymes was supported by both transcriptomics and proteomics (Dusfour et al., 2015; Epelboin et al., 2021; Faucon et al., 2017), those most contributing to deltamethrin resistance and the underlying genetic events remain to be identified. A recent study using a composite population from French Guiana resistant to multiple insecticides identified gene copy number variations (CNV) as a probable cause of detoxification enzymes overexpression in this region (Cattel et al., 2020). Through a pool-seq approach targeting > 300 candidate genes, this study identified multiple contiguous P450s from the CYP6 family on chromosome 1 showing an apparent elevated CNV in association with deltamethrin survival. However, despite the use of controlled crosses, the multigenic resistance phenotype of this population (carrying the three above-mentioned *Kdr* mutations at high frequency together with multiple metabolic resistance alleles) did not allow to conclude about their role in deltamethrin resistance. In addition, the exon-based nature of the sequencing data generated together with the complexity of *Ae. aegypti* genome did not allow resolving the genomic architecture of the duplicated locus.

In this context, the present study aimed at confirming the contribution of this P450 duplication in the resistance of *Ae. aegypti* to deltamethrin and at deciphering its genomic architecture. In order to limit confounding effects from other resistance alleles, two isofemale lines originating from French Guiana and showing contrasted pyrethroid resistance levels but no apparent resistance to organophosphate and carbamate insecticides were used (Epelboin et al., 2021). In addition, both lines lacked the two major *Kdr* mutations Val1016Ile and Val410Leu occurring in South America (Epelboin et al., 2021). Controlled crosses were then used to remove the Phe1534Cys *Kdr* mutation from the resistant line, which retained a P450-mediated resistance phenotype. Then, RNA-seq and whole genome pool-seq confirmed the presence of a ∼200 Kb genomic duplication enhancing the expression of a cluster of multiple P450 genes in the resistant line. The genomic architecture of the duplicated loci was elucidated through long read sequencing, providing clues about its evolutionary origin. Association studies, experimental evolution and reverse genetics were then used to further investigate the contribution of this duplicated allele to deltamethrin resistance.

## Results

### Deltamethrin resistance is associated with P450 activity and is autosomal

Two isofemale lines from Ile Royale island (French Guiana) showing contrasted resistance phenotypes, namely IR03 and IR13 lines, were used as starting material (**Table 1**). Comparative deltamethrin bioassays confirmed that the IR13 line can be categorized as susceptible to deltamethrin as previously shown (Epelboin et al., 2021), though its susceptibility was slightly lower than the laboratory strain Bora-Bora that was maintained for decades in insectaries. Mating IR03 genotyped individuals allowed producing the IR0F line deprived of the three *Kdr* mutations majorly associated with pyrethroid resistance in South America (*i*.*e*. Val410Leu, Val1016Ile, Phe1534Cys). However, our attempts to remove the only *Kdr* mutation remaining, Ile1011Met were not successful (see below). Bioassays showed that the IR0F line remained highly resistant to deltamethrin with only 44% mortality to a high dose of insecticide. Mortality levels remains similar between the IR03 and the IR0F lines suggesting that the removal of the Phe1534Cys *Kdr* mutation did not affect the resistance phenotype, or was compensated by the retaining of the Ile1011Met *Kdr* mutation.

**Table 1.** Origin, resistance status and *Kdr* mutations frequencies of the studied lines.

| Line | Origin, type | Deltamethrin mortality<sup>1</sup> | Resistance status | f(410Leu)<br>2 | f(1016Ile)<br>2 | f(1534Cys)<br>2 | f(1011Met)<br>3 |
| --- | --- | --- | --- | --- | --- | --- | --- |
| Bora-Bora | Fr. Polynesia, Lab | 100 | Susceptible | 0 | 0 | 0 | 0 |
| IR13 | Fr. Guiana, isofemale | 95 (91-99) | Susceptible | 0 | 0 | 0.26 | 0 |
| IR03 | Fr. Guiana, isofemale | 43 (26-60) | Resistant | 0 | 0 | 0.26 | 0.33 |
| IROF | IR03 cross | 44 (34-54) | Resistant | 0 | 0 | 0 | 0.50 |

<sup>1</sup> Mortality rates (mean % ± SD) previously obtained on 3 days-old females using 0.05% deltamethrin for 40 min (N = 10).
<sup>2</sup> *Kdr* mutation frequencies estimated by qPCR individual genotyping (N = 30) and later confirmed by pool-seq whole genome data.
<sup>3</sup> *Kdr* mutation frequencies estimated from pool-seq whole genome data.

The involvement of P450s in the resistance phenotype of the IR0F line was then confirmed by sequentially exposing adult mosquitoes to the P450 inhibitor piperonyl butoxide (PBO) and to the insecticide. Such PBO pre-exposure did not significantly affect deltamethrin survival in the susceptible IR13 line, but increased mortality from 2.2 % to 22.6 % in the IR0F line (Wilcoxon test P value = 0.009), supporting the contribution of P450-mediated detoxification in the resistance of the IR0F line (**Figure 1**).

**Figure 1.**
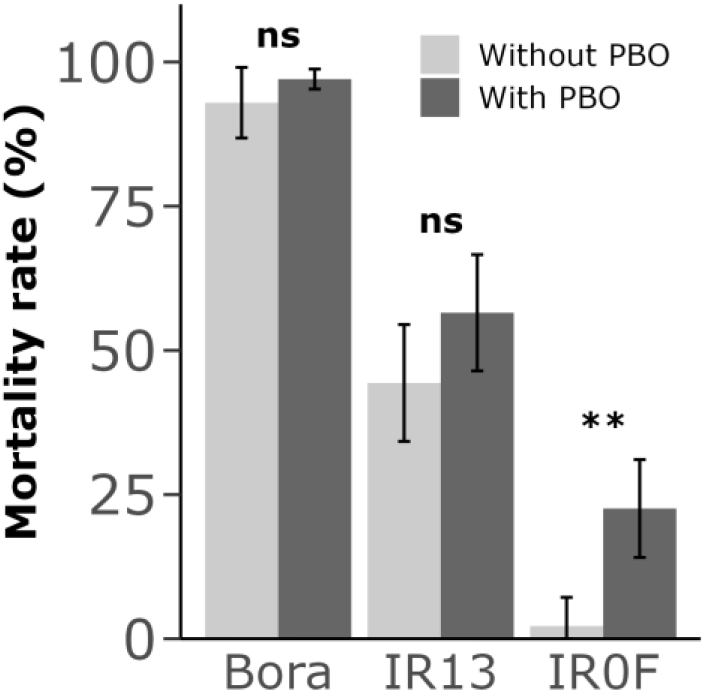
Synergistic effect of PBO on deltamethrin resistance. Adult females were exposed or not to 4 % PBO for 1h followed by a 30 min exposure to 0.03 % deltamethrin. Mortality was recorded 24h after exposure. Mean mortality rates are indicated ± SD, and compared using a Wilcoxon test (N = 5, ns: not significant; **: P value < 0.01).

As the duplicated P450 locus is located on chromosome 1, which also carries the sex determining locus, deltamethrin bioassays were performed on F1 and F2 males and females obtained from both ‘*Bora-Bora x IR0F’* reciprocal crosses in order to investigate the mode of transmission of resistance. Such comparative bioassays did not support any significant bias affecting the transmission of resistance (**Supplementary file 1**). For both F1 males and females, mortality levels upon deltamethrin exposure were similar in the two reciprocal crosses and intermediate to those of the parental lines, indicating the absence of a maternal effect and a semi-dominant phenotype in heterozygotes. Similar and intermediate mortality levels were again observed in F2 males and females, suggesting the lack of a sex transmission bias.

### Overexpression of the duplicated CYP6 gene cluster

RNA-seq was used to identify differentially transcribed genes between the two resistant lines (IR03 and IR0F), and the two susceptible lines (IR13 and Bora-Bora). Among the 11268 protein-coding genes passing our coverage filter (82 % of all protein coding genes), 84 genes showed a significant and consistent differential transcription level in the four ‘*resistant vs susceptible’* pairwise comparisons (see methods, expression fold change ≥ 1.5 and adjusted P value ≤ 0.0005). The 39 genes under-transcribed in resistant lines did not include any gene belonging to any family known to be associated with insecticide resistance. Conversely, the 45 genes over-transcribed in resistant lines included eight genes belonging to gene families associated with metabolic resistance (**Figure 2** and **Supplementary file 2**). These included five P450s (*CYP6BB2* AAEL014893, *CYP6*-*like* AAEL026852, *CYP6-like* AAEL017061, *CYP6P12* AAEL014891 and *CYP9F-like* AAEL028635). The four *CYP6* genes showed mean fold changes up to 4.0 fold and belong to a cluster of five consecutive *CYP6* genes located at ∼ 271 Mb on chromosome 1. Among them, *CYP6P12* (AAEL014891) was annotated as a single accession in the reference genome while both protein sequence and RNA-seq data support the presence of two distinct genes hereafter referred to as *CYP6P12v1* and *CYP6P12v2* when appropriate.

**Figure 2.**
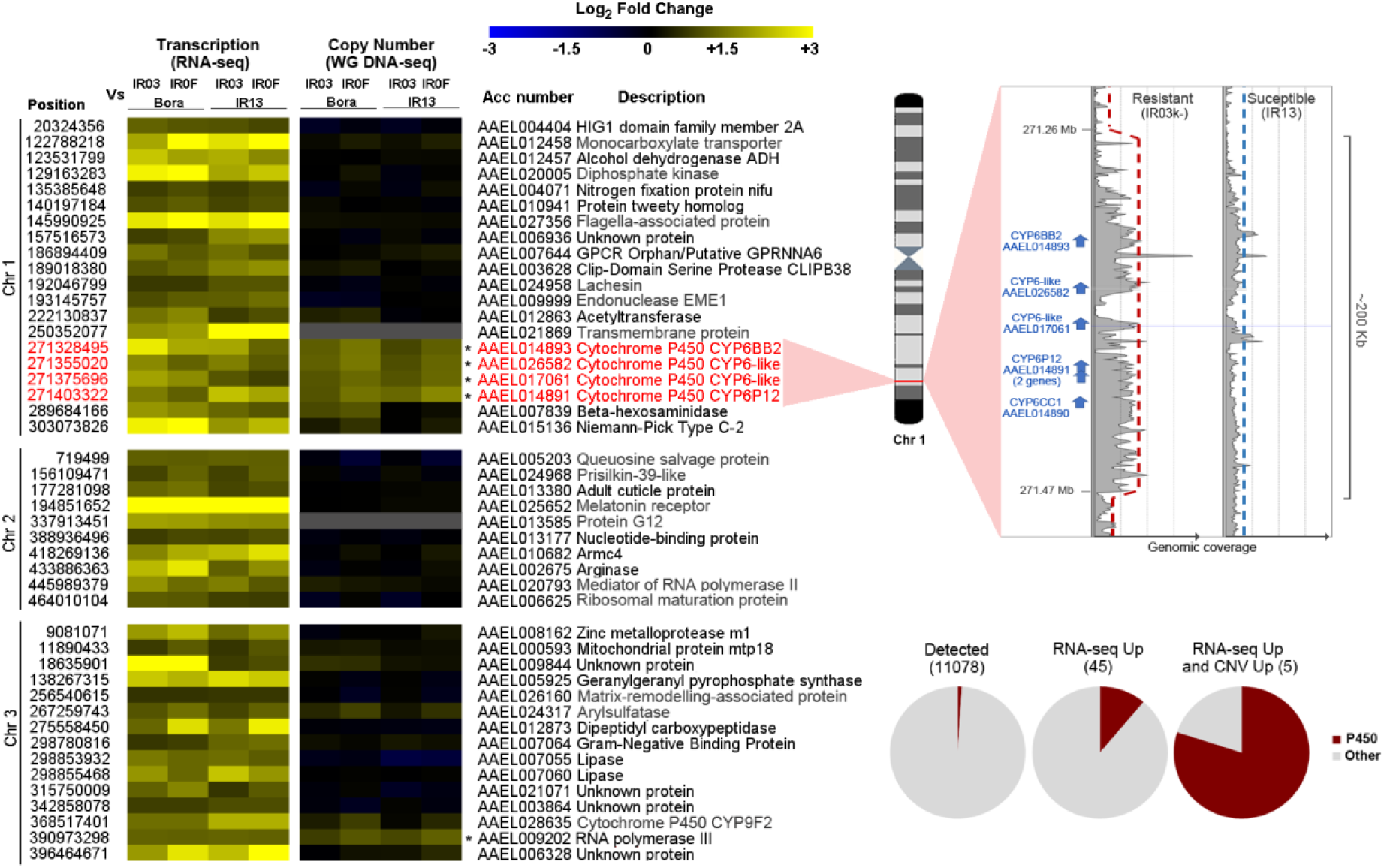
Genes over transcribed in resistant lines and their associated copy number variations. Only the 45 genes found significantly over transcribed from RNA-seq are shown (FC ≥ 1.5 and corrected P value ≤ 0.0005 in all resistant versus susceptible pairwise comparisons). Genes also showing an elevated copy number from WG DNA-seq (FC ≥ 1.5) in all pairwise comparisons are indicated with stars. Genomic coordinates (chromosome and start position), accession number and description are indicated for all genes. Gene descriptions in grey were obtained from a BlastP against NCBI Refseq. Genes in red belong to a single duplicated region on chromosome 1. Pool-seq read coverage profiles of the susceptible IR13 and the resistant IR0F lines around the duplicated locus are shown. Pie charts show the proportion of P450s in each dataset.

Whole genome pool-seq short read data were then used to identify genes affected by copy number variations in association with resistance base on their normalised exonic coverage. This analysis identified 22 genes showing a congruent elevated copy number (CN) in resistant *vs* susceptible lines (FC ≥ 1.5 in all *‘resistant vs susceptible’* pairwise comparisons). These included seven P450s. One of them (AAEL009018) was located at ∼ 58.8 Mb on chromosome 1 and was not detected as over-transcribed from RNA-seq. The six others were located at ∼271 Mb on chromosome 1 and included the five *CYP6s* found over-transcribed in resistant lines (see above) and another *CYP6* (*CYP6CC1*) from the same gene cluster that was filtered out from RNA-seq analysis due to its low transcription level (**Figure 2** and **Supplementary file 2**). The over-transcribed CYP9F-like located on chromosome 3 showed a slight increased CN in the IR0F line *vs* the Bora-Bora line (1.6 fold) but such increased CN was not confirmed for the three other ‘resistant *vs* susceptible comparisons’ (1.07, 1.4 and 1.2 fold for IR03/IR13, IR0F/IR13 and IR03/Bora respectively). A closer look at the genomic coverage at the CYP6 locus on chromosome 1 revealed a ∼ 2 fold increased coverage affecting a region of ∼ 200 Kb spanning the entire CYP6 cluster. Such duplication was observed in all resistant *vs* susceptible comparisons. Altogether, these genomic data showed that the deltamethrin resistance phenotype observed in the resistant lines is associated with the presence of a large duplication on chromosome 1 affecting the transcription level of six clustered *CYP6* genes.

Whole genome short reads data confirmed the absence of the three major *Kdr* mutations Val410Leu, Val1016Ile and Phe1534Cys in the IR0F resistant line, while another *Kdr* mutation (Ile1011Met) was identified at an exact 0.5 frequency (141/282 short reads supporting the 1011Met allele). Indeed, both short read and long read data confirmed that this Ile1011Met *Kdr* mutation is affected by an heterogenous genomic duplication (see below and in Martins et al., 2013). Therefore, a 0.5 frequency indicates that the Ile-Met/Ile-Met genotype is fixed in the IR0F line.

### Genomic architecture of the duplicated loci

Though not the major aim of the present study, our genomic data allowed characterizing the architecture of the duplication affecting the Ile1011Met *Kdr* mutation in the IR0F line. Both short read and long read topologies confirmed that the Ile1011Met *Kdr* mutation is part of 125 Kb genomic duplication covering the 21 last coding exons of the VGSC gene AAEL023266 (**Supplementary file 3**). The breakpoints of this partial gene duplication are located at 315,905,811 bp and 316,030,250 bp on the reference genome AaegL5. The partial 3’ copy carries the wild type allele (1011Ile) while the full-length copy carries the resistant allele (1011Met). The two copies also differed by their intronic sequence with the partial copy bearing a B type intron and the full-length copy bearing a A type intron as previously described (Martins et al., 2013). As expected from an incomplete gene duplication, RNA-seq data supported the sole expression of the 1011Met allele meaning that the IR0F line (genotype Ile-Met/Ile-Met) expresses a Met phenotype.

The genomic structure of the P450 duplicated region identified on chromosome 1 was further investigated in light of long reads data obtained from the IR13 susceptible line and the IR0F resistant line. This revealed the presence of a 220.4 Kb duplication in the IR0F line, with the two tandem repeats separated by a 6 Kb insertion (**Figure 3**). The right breakpoint of the duplication (RB) was found ∼ 50 Kb right to *CYP6CC1* at position 271,472,082 bp, and the left breakpoint (LB) ∼ 77 Kb left to *CYP6BB2* at 271,251,705 bp. These breakpoints were supported by split reads from both long and short read data (**Supplementary file 4**). The duplication is flanked by two distinct transposons, each having its own terminal inverted repeats (TIR): a 2.4 Kb PiggyBac-like transposon on the right (PYL), and 9 Kb hAT-related transposon on the left (hAT-CYP6). These transposons are described in more detail in **Supplementary file 5**.

**Figure 3.**
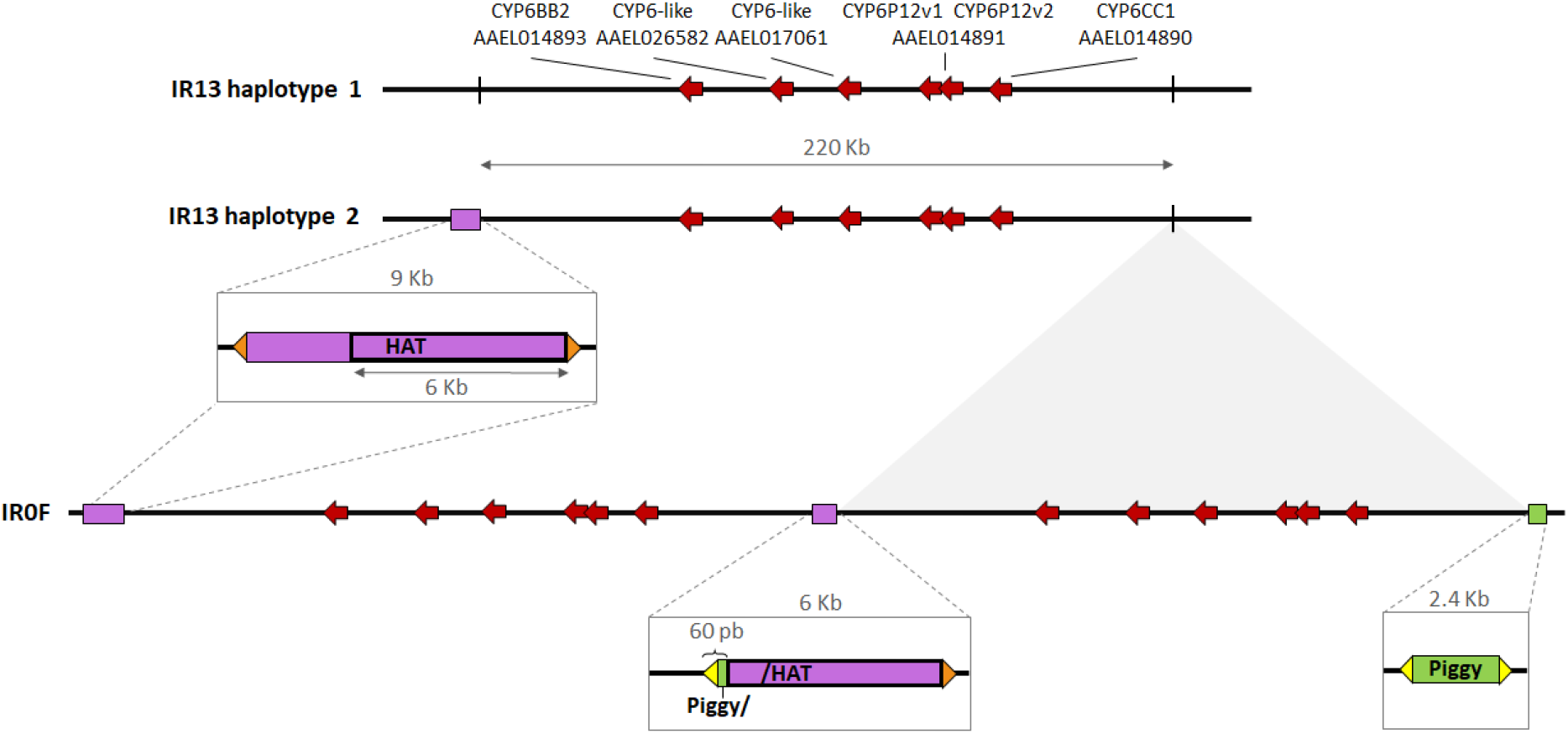
Genomic architecture of the duplication. The topology was deduced from both short and long reads sequencing data. The six CYP6 genes carried by the duplication in the resistant lines are represented by red arrows. Regions containing full or partial transposable elements are shown in greater detail. Orange and yellow triangles denote the inverted terminal repeats from hAT (17 pb) and PiggyBac-like (18 pb) elements respectively. Diagram is not to scale.

The central sequence joining the tandem copies is chimeric, with 60 bp from the left end of PYL transposon followed by 6001 bp from the right end of the hAT transposon. This duplication is absent from the IR13 susceptible line (no read joining RB and LB sequences and no PYL transposon sequence, neither full nor partial at RB). However, the hAT-CYP6 transposon was identified at the LB in one IR13 haplotype, much like in the IR0F line, while a second haplotype was devoid of hAT insertion, much like the reference genome. An excess of tri-allelic SNPs is expected in the duplicated region if the two copies have diverged. Pool-seq short read data were then used to compare the frequency of tri-allelic loci within the duplication and 100 Kb upward and downward. Among the 1401 substitutions sites identified within the duplication, five were tri-allelic whereas no such tri-allelic variant was identified among the 1334 substitutions identified in the flanking regions. This represents a slight but significant enrichment in the duplicated region (Fisher test P value = 0.031). These tri-allelic sites were all located within a 11.2 Kb intergenic region between the two *CYP6*-*like* genes AAEL017061 and AAEL014891v1.

### The P450 duplication and the Kdr 1011 mutation are both associated with deltamethrin survival

The association of the P450 duplication and the *Kdr* Ile1011Met mutation with deltamethrin survival were investigated in F2 individuals obtained from both ‘Bora-Bora x IR0F’ reciprocal crosses. F2 females were then exposed to a high dose of deltamethrin (85.5 % mortality) before quantifying *CYP6BB2* gene copy number and the frequency of the 1011Met *Kdr* allele in dead and survivors using ddPCR (**Figure 4**). As expected in the presence of a fixed duplication, a 2 fold increase of *CYP6BB2* copy number was observed in the IR0F line (4.31 ± 0.21 copies) as compared to Bora-Bora line (2.07 ± 0.10 copies) while F1 individuals showed an intermediate copy number (3.58 ± 0.14 copies). In F2 individuals, *CYP6BB2* copy number was higher in survivors than in dead individuals (4.00 ± 0.16 copies versus 3.31 ± 0.15 copies) supporting a genetic linkage between the P450 duplication and deltamethrin survival. Insecticide survival was also linked to the *Kdr* mutation Ile1011Met on chromosome 3. In line with an heterogenous *Kdr* duplication, the frequency of the 1011Met allele was close to 0.5 in the IR0F resistant line (genotype Ile-Met/Ile-Met) and close to 0.33 in F1 individuals (genotype Ile-Met/Ile). In deltamethrin-exposed F2 individuals, the Met allele frequency was higher in survivors than in dead individuals (0.41 ± 0.02 versus 0.30 ± 0.02).

**Figure 4.**
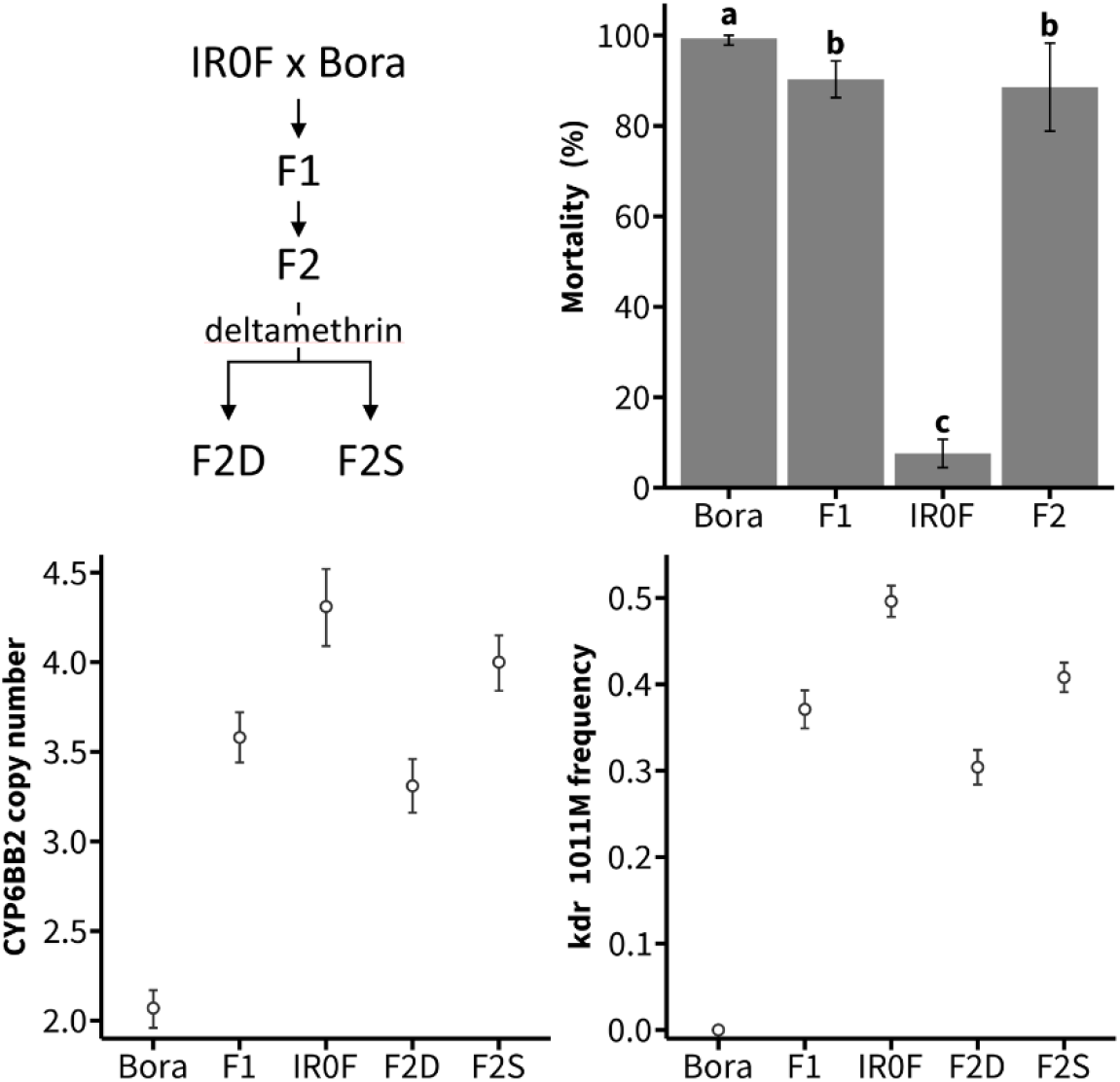
Genotype-phenotype association study. Top left panel shows the crossings underlying this experiment. Top right panel shows mortality data obtained by exposing adult females from the IR0F and Bora-Bora lines and their F1 and F2 progeny to 0.03 % deltamethrin for 1 hour. Mean mortalities ± SD 24 h after insecticide exposure are shown and distinct letters indicate significant differences (Kruskal-Wallis test followed by post hoc Wilcoxon test with Bonferroni-Holm correction, N ≥ 5, p ≤ 0.05). Bottom left and right panels show the copy number of *CYP6BB2* and the frequency of the *Kdr* 1011Met allele respectively as inferred by ddPCR from pools of individuals (mean ± 95 % CI, Bora: N = 19, IR0F: N = 23, F1: N = 17, F2D: N = 113, F2S: N = 15). Dead and surviving F2 individuals 24 h after insecticide exposure are noted F2D and F2S respectively.

### Multiple CYP genes carried by the P450 duplication may contribute to resistance

The relative importance of the P450s carried by the duplication in deltamethrin survival was then examined by RNA interference. Direct dsRNA injections in adult females of the IR0F line were used to specifically knock down the five *CYP6* genes showing sufficient expression to be detected by RNA-seq (*CYP6BB2, CYP6*-*like* AAEL026582, *CYP6-like* AAEL017061, *CYP6P12v1* and *CYP6P12v2*). Such approach allowed reaching an acceptable silencing specificity and a moderate silencing efficiency (from 42.2 % for AAEL017061 to 69.7 % for AAEL026582). Comparative deltamethrin bioassays performed on dsCYP6-injected mosquitoes and dsGFP-injected controls suggested that the two first CYP genes of the duplicated cluster (*CYP6BB2* and *CYP6-like* AAEL026582) contribute to deltamethrin survival (see methods and results in **Supplementary file 6**). However, data generated across multiple injection experiments were not fully conclusive because a high mortality was frequently observed in dsGFP-injected controls and because the increased mortalities observed in dsCYP-injected mosquitoes were relatively low. Such low mortality variations upon dsCYP6 injection were likely due to the moderate silencing efficiency and the presence of the *Kdr* 1011Met allele in the IR0F line.

### The P450 duplication is hardly retained by selection in presence of the Kdr 1011 allele

The co-occurence of the P450 duplication and the Ile1011Met *Kdr* mutation in the IR0F resistant line was taken as an opportunity to compare their responses to deltamethrin selection. Four independent lines obtained from ‘Bora-Bora x IR0F’ F3 offspring and representing two replicated selection regimes (selected lines: SelA and SelB, non-selected lines: NSA and NSB) were compared for their resistance level, *CYP6BB2* copy number and *Kdr* 1011Met allele frequency (**Figure 5**).

**Figure 5:**
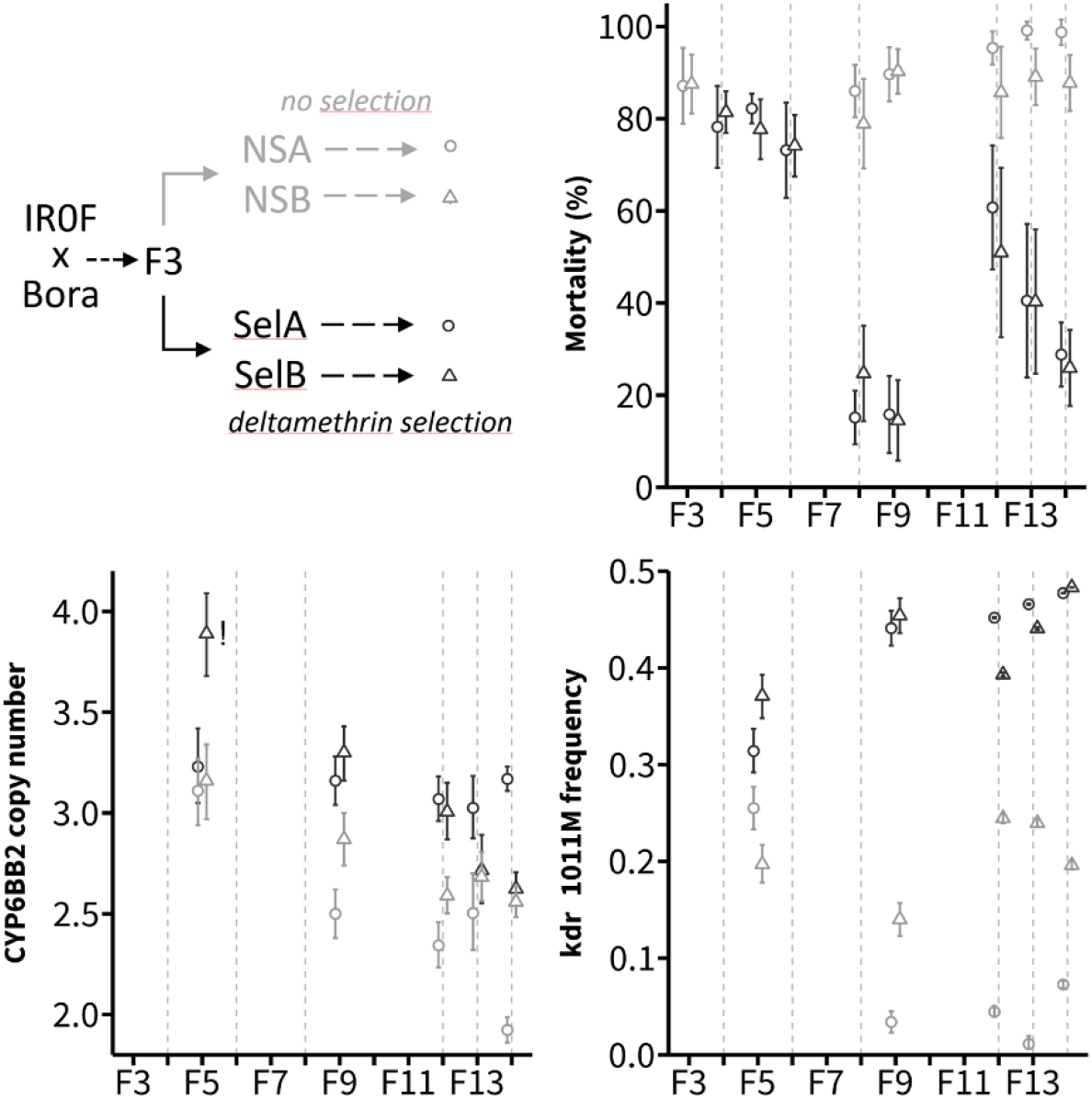
Response of the P450 duplication and the *Kdr* 1011Met allele to deltamethrin selection. Top left panel shows the crossings underlying this experiment. Top right panel shows the resistance of each line as measured by bioassays on adult females (0.03 % deltamethrin for 1h exposure, mean mortality ± SD). Bottom left and right panels show CYP6BB2 copy number and *Kdr* 1011Met frequency respectively as inferred by ddPCR from pools of individuals (mean ± 95 % CI, N ≥ 30). The SelB *CYP6BB2* copy number data point marked with a ‘!’ was caused by an unexpected sampling error and was therefore excluded from the statistical analysis comparing *CYP6BB2* copy number between Sel and NS lines across generations. Grey dashed lines indicate the generations for which Sel lines were selected with deltamethrin.

An increase of deltamethrin resistance was observed for both Sel lines, with mortality dropping from ∼80 % in F4 to less than 20 % in F9. Relaxing deltamethrin selection for three successive generations (F9 to F11) led to an increased mortality in both Sel lines, suggesting that the resistance phenotype is costly. Though subjected to sampling effects, ddPCR data showed that the *Kdr* 1011Met allele was rapidly selected by deltamethrin. Its frequency was close to 0.5 in F9 of both Sel lines, supporting the near fixation of the Ile-Met duplicated allele. Such increased frequency was not seen in NS lines where Met allele frequency decreased to less than 10% in one line and fluctuated under 0.25 in the other line. No such strong response to selection was observed for the P450 duplication, with *CYP6BB2* copy number showing a gradual decrease through generations in both Sel and NS lines. However, *CYP6BB2* mean copy number was significantly higher in Sel lines than in NS lines across generations, supporting a higher retention rate of the duplicated allele upon deltamethrin selection (Linear Mixed Effect Model, P value = 0.0088).

## Discussion

In line with their high probability of occurrence, genomic duplications have been shown to frequently contribute to short-term adaptation in various organisms (Kondrashov, 2012). In arthropods facing strong selection pressures from insecticides, duplications are known to be adaptive via two distinct mechanisms (Bass & Field, 2011). First, heterogeneous duplications affecting key insecticide target genes and tethering both susceptible and resistant alleles have been associated with a reduction of fitness costs (Labbe et al., 2007). Second, duplications of detoxification genes can contribute quantitatively to their over-expression, leading to resistance through increased insecticide metabolism (Faucon et al., 2015; Weetman et al., 2018). As opposed to target-site resistance mutations that can be easily tracked by PCR in natural populations, the paucity of DNA markers underlying metabolic resistance hinders their tracking in the field. In this concern, identifying gene duplications associated with metabolic resistance can provide new tools to monitor the dynamics of resistance alleles (Cattel et al., 2021).

### The P450 duplication affects a cluster of genes previously associated with resistance

The present study confirmed the occurrence of a large genomic duplication affecting a cluster of six P450s from the *CYP6* family in *Ae. aegypti* populations from French Guiana. The presence of such a duplication was previously suspected though the structure of the duplicated loci could not be resolved from targeted-sequencing data (Faucon et al., 2015). The association of this P450 duplication with deltamethrin resistance was supported by controlled crosses, although the presence of other resistance alleles limited the association power (Cattel et al., 2020). A duplication encompassing the *CYP6BB2* gene was also associated with pyrethroid resistance in Lao PDR (Marcombe et al., 2019). Though gene duplication was not looked for, the over-transcription of CYP6 genes from this cluster (e.g. *CYP6BB2, CYP6P12* and *CYP6CC1*) was frequently associated with pyrethroid resistance worldwide (Bariami et al., 2012; Dusfour et al., 2015; Goindin et al., 2017; Kasai et al., 2014; Marcombe, Poupardin, et al., 2009; Moyes et al., 2017; Reid et al., 2014; Saavedra-Rodriguez et al., 2012; Seixas et al., 2017). These genes were also found under directional selection in association with pyrethroid survival in a Mexican resistant line from whole exome SNP data (Saavedra-Rodriguez et al., 2021). In addition, *CYP6BB2* was also shown capable of metabolising the pyrethroid permethrin (Kasai et al., 2014). By comparing its copy number and transcriptional level across different lines, Faucon et al., (2017) suggested that both transcriptional regulation and genomic duplication contribute to its over-expression in South America. Interestingly, this gene also responded to selection with the neonicotinoid imidacloprid, suggesting that P450s from this cluster can be selected by and metabolise other insecticides (Riaz et al., 2013; Zoh et al., 2021).

Our attempt to isolate this P450 duplication from other resistance alleles occurring in French Guiana was partially successful. Indeed, both isofemale lines were naturally deprived from the major Val1016Iso and Val410Leu *Kdr* mutations, and controlled crosses allowed removing the Phe1534Cys mutation. However, the resulting IR0F resistant line carrying the P450 duplication still carried the Ile1011Met *Kdr* mutation as a fixed heterogenous duplication (Ile-Met/Ile-Met genotype, Met/Met phenotype). There is evidence that this mutation confers some resistance to pyrethroids in the field (Brengues et al., 2003; Brito et al., 2018; Martins et al., 2009). Nevertheless, the use of the P450 inhibitor PBO showed that the resistance phenotype was still associated with P450-mediated metabolism. Considering that four of the five P450s over-transcribed in resistant lines belong to the duplicated locus, the contribution of this P450 duplication to the resistance phenotype is likely. Studying the mode of transmission of resistance did not allow evidencing any significant sexual transmission bias potentially associated with resistance. In line with this, this P450 duplication is located outside of a 210 Mb region of chromosome 1 showing a low recombination rate and a high genetic differentiation between *Ae. aegypti* males and females (Fontaine et al., 2017).

### Contrasted architectures of the duplications affecting the P450 and the Kdr loci

The presence of two distinct transposons at the P450 duplication breakpoints implies that they have played an active role in the generation of this genomic event. Transposon-mediated duplication often arises from ectopic recombination between two homologous transposon copies inserted in the same orientation at distinct positions of a same chromosome (Baker et al., 1996). However, such an event would leave a single full-length transposon copy between the two duplicated genomic regions with no transposon at other breakpoints (Remnant et al., 2013). Here, a more complex scenario must have taken place because two distinct transposons are flanking the duplication, and because the sequence joining the two copies is chimeric. A possible scenario involves independent insertion of the two distinct transposons, then transposition of the hAT into the PiggyBac-like transposon followed by a deletion event, and finally an ectopic homologous recombination event between the two duplicated hAT sequences (**Supplementary File 7**). One transposon involved is related to the hAT superfamily while the other one is distantly related to PiggyBac (see **Supplementary file 5** for more details), both being cut and paste elements having multiple copies and being active in the *Ae. aegypti* genome (Nene et al., 2007). The divergence of the two tandem copies was supported by the identification of a few tri-allelic substitution sites within the duplicated region, most likely implying mutations occurring post-duplication. The few tri-allelic sites observed represent only a fraction of those, because they imply a pre-duplication mutation at the same position, and because mutations reverting to the reference allele are not detected as such. Nevertheless, such low divergence rate between the two copies supports the recent age of this duplication event.

In contrast, the genomic duplication affecting the VGSC gene at the *Kdr* Ile1011Met locus shows no remnant of any transposon at breakpoints, suggesting a different genesis than the P450 duplication. It is likely to be an earlier event, since the copy divergence is much more pronounced, particularly in intronic regions (Martins et al., 2013). Also, it differs from other heterogenous duplications implicated so far in insecticide resistance because the incomplete copy carrying the susceptible Ile1011 pseudoallele, does not contribute to gene expression. It therefore cannot modulate the resistance level provided by the 1011Met allele carried by the full copy, nor its fitness. This contrasts with the heterogenous duplications affecting the *Ace1* target-site mutation observed in *Culex pipiens sp*. and *Anopheles gambiae*, where both copies can contribute to the phenotype, thus allowing a reduction of the fitness cost associated with the resistant allele in absence of insecticide (Djogbenou et al., 2009; Labbe et al., 2007). In this regard, further investigating the evolutionary origin and dynamics of this *kdr* heterogenous duplication deserves further attention.

### The P450 duplication has a limited adaptive value in presence of the Kdr 1011 mutation

The association study performed on F2 individuals supported the association of the P450 duplication with deltamethrin survival together with the Ile1011Met *Kdr* mutation. The association of the Ile1011Met *Kdr* mutation with deltamethrin survival was expected. As the *VGSC* gene carrying *Kdr* mutations is located on chromosome 3, a genetic linkage with the P450 duplicated loci is excluded. Therefore, the association of the P450 duplication in F2, following limited recombination events, implies a resistance locus located on chromosome 1, either the P450 duplication itself or another genetically linked locus. However, no other known resistance gene was identified in this genomic region. The relative importance of the different *CYP6* genes carried by the P450 duplication in deltamethrin survival was investigated through RNA interference. Despite a limited statistical power, likely due to the presence of the *Kdr* Ile1011Met resistance allele and moderate knock down efficiencies, these data suggest that at least two *CYP6* genes (*CYP6BB2* and AAEL026582) carried by the duplication may contribute to deltamethrin detoxification.

Experimental evolution confirmed the strong positive response of the *Kdr* 1011Met allele to selection with a high dose of deltamethrin. Such experiment also evidenced a gradual decrease of this allele in absence of insecticide selection, supporting a significant fitness cost as previously shown for other *Kdr* mutations (Rigby et al., 2020; Uemura et al., 2023). The response of the P450 gene duplication to deltamethrin selection was less clear. A gradual decrease of *CYP6BB2* copy number was observed in both selected and non-selected lines, even though such decay was slower in the selected lines. Such result supports a lower adaptive value of the P450 duplication in our experimental conditions. Such unexpected response may indicate the presence of a significant fitness cost associated with this large genomic duplication. Indeed, genomic duplications can be detrimental for various reasons (Schrider et al., 2013): first, gene duplications can alter the gene dosage balance which can lead to metabolic cost. Second, genomic duplications allow the accumulation of deleterious mutations that are less cleared off by background selection because of the functional redundancy of duplicated copies. Third, large duplications can impair recombination locally. In the present case, metabolic costs due to gene dosage balance and/or a low recombination rate at this locus between the Bora-Bora line (well adapted to laboratory conditions) and the IR0F line (less fitted to our laboratory conditions) may have impaired the selection of the P450 duplication and favoured the selection of other resistance alleles such as the *Kdr* 1011Met mutation. Finally, considering the fitness costs associated with this *Kdr* mutation, its additive cost with the P450 duplication may have prevented their concomitant selection with the use of a high insecticide dose favouring the selection of the former. Such fitness-cost balance at two distinct resistance loci may reflect the complex interactions occurring between resistance alleles *in natura*, leading to distinct adaptive trajectories depending on selection pressures and demographic effects.

### Conclusions

Following previous work, the present study supports the contribution of this P450 gene duplication in pyrethroid resistance. However, deciphering its adaptive value *versus* other resistance alleles and its dynamics in natural mosquito populations deserves further work. The relative importance of the different *CYP6* genes carried by this duplication in resistance to pyrethroids (and possibly to other insecticides) also deserves further validation. Despite this incomplete picture, the nature of the P450s affected by this large genomic duplication and their frequent association with insecticide resistance call for further studying its evolutionary origin and dynamics in response to xenobiotics. In this context, though the flanking of the duplication by repeated transposable elements prevented the design of a specific PCR diagnostic assay targeting the breakpoints, the dual-colour quantitative TaqMan ddPCR assay we developed (see **Supplementary file 8**) represents a good alternative to track this resistance allele in natural mosquito populations.

## Methods

### Mosquitoes

The *Ae. aegypti* laboratory strain Bora-Bora, fully susceptible to insecticides was used as reference in the present study. The isofemale lines used in the present study were derived from a field population collected in 2015 in the Ile Royale (IR) island (5.287° N; 52.590° W) off the coast of French Guiana as described in Epelboin et al., (2021). Among the isofemale lines isolated from this population, the lines IR13 and IR03 showed contrasted pyrethroid resistance levels with the IR03 line being highly resistant and the IR13 line being susceptible as defined by WHO diagnostic dose bioassay (Table 1) and Epelboin et al., (2021). Both lines were shown to lack the two major voltage-gated sodium channel *Kdr* mutations occurring in this geographical area (Val410Leu and Val1016Ile), yet both carried the Phe1534Cys mutation at a moderate frequency (Epelboin et al., 2021). The proteomic profiling of these lines identified multiple cytochrome P450s enriched in the IR03 resistant line as compared to a susceptible line, supporting the presence of metabolic resistance alleles (Epelboin et al., 2021). All mosquito lines were maintained under standard laboratory conditions (27 ± 2°C, 70 ± 10 % relative humidity, light/dark cycle 14:10 h) and large population size to limit drift effects (N > 1000). Larvae were reared in tap water and fed with hay pellets. Adults were maintained in mesh cages and fed with a 10 % honey solution. Blood feedings were performed on mice.

### Removal of the Phe1534Cys mutation from the IR03 line

In an attempt to isolate metabolic resistance alleles, inter-crosses assisted by genotyping were performed on the IR03 resistant line to create the IR0F line depleted from the Phe1534Cys *Kdr* mutation. A total of 245 IR03 couples were individualised at the pupal stage in Eppendorf tubes placed in plastic cups covered by a nylon mesh and allowed to emerge and reproduce. The Phe1534Cys *Kdr* mutation was first searched for from male exuvia for all couples and then from female exuvia of the 58 couples from which males were negatives for the 1534Cys allele. Genotyping was performed on qPCR using the High Resolution Melt Curve Analysis method as described in Saavedra-Rodriguez et al., (2007). The six couples in which both parents were homozygous wild type (Phe-Phe) were transferred to a mesh cage, blood fed and allowed to lay eggs. The absence of the 1534Cys allele in the offspring was confirmed by genotyping 40 individuals of both sexes. Adults were then exposed to 0.05 % deltamethrin and survivors were allowed to freely reproduce to generate the IR0F resistant line. The IR0F line was then maintained in standard insectary conditions under moderate deltamethrin selection pressure (*i*.*e*. 50 % mortality every two generations).

### Deltamethrin bioassays

All bioassays were performed by exposing replicates of 20-25 three days-old non-blood-fed adults to deltamethrin impregnated filter papers following the standard WHO procedure, with mortality being recorded 24h after insecticide exposure (WHO, 2022). Different insecticide doses and exposure times were used according to the nature of the experiment and the mosquito tested (susceptibility status, sex, use of enzymatic inhibitor or not) in order to keep mortality in the dynamic range and maximise mortality variations between conditions. The bioassays used for assessing the resistance level of the different lines presented in Table 1 were performed on females using 10 replicates per line and a dose of 0.05% deltamethrin for 40 min. The involvement of P450s in the resistance phenotype presented in Figure 1 was assessed by bioassays on 5 replicates of females exposed or not to 4 % piperonyl butoxide (PBO) for 1 h prior to deltamethrin exposure using a dose of 0.03 % deltamethrin for 30 min. Bioassays used to investigate the mode of inheritance of resistance presented in Supplementary Figure 1 were performed on males and females obtained from the susceptible Bora-Bora line and the resistant IR0F line together with F1 and F2 individuals obtained from each reciprocal crosses (see below). As males and females naturally show different tolerance to insecticides and F1/F2 individuals were expected to be more susceptible than IR0F individuals, deltamethrin exposure conditions were accordingly (0.03% deltamethrin for 30 min for females and 0.015% deltamethrin for 25 min). Finally, bioassays used for monitoring the resistance level of the selected and non-selected lines through the experimental evolution experiment presented in Figure 5 were performed on 5 to 7 replicates per line per generation using a dose of 0.03 % deltamethrin for 1 h.

### Controlled crosses and association studies

In order to investigate the mode of inheritance of resistance, reciprocal crosses between the fully susceptible Bora-Bora line and the resistant IR0F line were performed (cross A: ‘Bora-Bora females x IR0F males’ and cross B: ‘IR0F females x Bora-Bora males’). Crosses were performed with 200 virgin females and males of each line. The susceptibility of F1 and F2 males and females from each cross were then compared using deltamethrin bioassays as described above. The association of the P450 duplication with deltamethrin survival was studied in F2 females obtained from both reciprocal crosses pooled in equal quantities. F2 females were exposed to 0.03 % deltamethrin for 1 h leading to 88.5 % mortality after a 24 h recovery time. Dead and surviving females were sampled and stored at -20°C until molecular analyses. The response of the P450 duplication and the Ile1011Met *Kdr* mutation were further studied across multiple generations using experimental selection. F3 eggs obtained from both reciprocal crosses were pooled in equal quantity and then randomly split in 4 lines: the first two lines (NSA and NSB) were maintained without insecticide selection, while the two other lines (SelA and SelB) were selected with deltamethrin. This duplicated line setup was used in order to control for genetic drift that may occur across generations. Deltamethrin mass selection was performed prior mating on both virgin males and virgin females (N > 1000 for each line) at generations F4, F6, F8, F12 and F13. A constant dose of deltamethrin was used through the selection process leading to ∼ 70 % mortality in each sex at generation F3 (females: 0.03 % for 1 h; males: 0.015 % for 30 min). The resistance level of each line was monitored at generations F5, F9, F12, F13 and F14 on females prior to insecticide selection using standard bioassays (see above). Unexposed adult females from each line were sampled at the same generations and stored at -20 °C until molecular analysis.

### RNA-sequencing

Gene transcription levels were compared across the four lines (Bora-Bora, IR13, IR03 and IR0F) using RNA-seq. For each line, four RNA-seq libraries were prepared from distinct batches of 25 calibrated three-day-old non-blood-fed females not exposed to insecticide. Total RNA was extracted using TRIzol (Thermo Fisher Scientific) following manufacturer instructions. RNA samples were then treated with RNase-free DNase set (Qiagen) to remove gDNA contamination, purified on RNeasy mini columns (Qiagen) and QC checked using Qubit (Thermo Fisher Scientific) and bioanalyzer (Agilent). RNA-seq libraries were prepared from 500 ng total RNA using the NEBNext® Ultra™ II directional RNA library Prep Kit for Illumina (New England Biolabs) following manufacturer’s instructions. Briefly, mRNAs were captured using oligodT magnetic beads and fragmented before being reverse transcribed using random primers. Double-stranded cDNAs were synthesised, end-repaired, and adaptors were incorporated at both ends. Libraries were then amplified by PCR for 10 cycles and purified before QC check using Qubit and Bioanalyzer. Libraries were then sequenced in multiplex as single 75 bp reads using a NextSeq500 sequencer (Illumina). An average of 51.4 M reads was generated per library. After unplexing and removing adaptors, sequenced reads from each library were loaded into Strand NGS V3.2 (Strand Life Science) and mapped to the latest *Ae. aegypti* genome assembly (Aaeg L5) using the following parameters: min identity = 90 %, max gaps = 5 %, min aligned read length = 25, ignore reads with > 5 matches, 3’ end read trimming if quality < 20, mismatch penalty = 4, gap opening penalty = 6, gap extension penalty = 1. Mapped reads were then filtered based on their quality and alignment score as follows: mean read quality > 25, max N allowed per read = 5, alignment score ≥ 90, mapping quality ≥ 120, no multiple match allowed, read length ≥ 35. These filtering steps allowed retaining ∼ 75 % of reads. Quantification of transcription levels was performed on the 13614 protein-coding genes using the DESeq2 method with 1000 iterations (Anders & Huber, 2010, https://bioconductor.org/packages/release/bioc/html/DESeq2.html). Only the 11268 protein-coding genes showing a normalised expression level ≥ 0.5 (∼ 0.05 RPKM) in all replicates for all lines were retained for further analysis. Differential gene transcription levels between each line across all replicates were then computed using a one-way ANOVA followed by a Tukey post hoc test and P values were corrected using the Benjamini and Hochberg multiple testing correction (Benjamini & Hochberg, 1995). Genes showing a transcription variation ≥ 1.5 fold (in either direction) and an adjusted P value ≤ 0.0005 in the four pairwise comparisons between resistant and susceptible lines (*i*.*e*. IR03 *vs* Bora-Bora, IR03 *vs* IR13, IR0F *vs* Bora-Bora and IR0F *vs* IR13), were considered differentially transcribed in association with deltamethrin resistance.

### Whole genome pool-sequencing

The genome of the four lines were compared using short read whole genome pool-seq. For each line, genomic DNA was extracted from two batches of 50 adult females using the PureGene Core Kit A (Qiagen) and the two gDNA extracts were then pooled in equal proportion into a single sequencing library. Whole genome sequencing was performed from 200 ng gDNA. Sequencing libraries were prepared according to the TruSeq DNA Nano Reference guide for Illumina Paired-end Indexed sequencing (version Oct. 2017) with an insert size of 550 bp. Sequencing was performed on a NextSeq 550 (Illumina) as 150 bp paired-reads. Sequencing depth was adjusted to reach an average coverage ≥ 80 X leading to the sequencing of ∼713 M reads per library. Reads obtained from each line were loaded into Strand NGS V3.2 and mapped to *Ae. aegypti* genome (Aaeg L5) using the following parameters: min identity = 90 %, max gaps = 5 %, min aligned read length = 25, ignore reads with > 5 matches, 3′ end read trimming if quality < 15, mismatch penalty = 4, gap opening penalty = 6, gap extension penalty = 1. Mapped reads were then filtered based on their quality and alignment score as follows: mean read quality > 25, max N allowed per read = 5, alignment score ≥ 90, mapping quality ≥ 60, no multiple match allowed, read length ≥ 100. Finally, interchromosomal split reads and PCR duplicates were removed. These filtering steps allowed retaining ∼53 % of reads.

In order to focus on duplication impacting protein expression, gene copy number variation (CNV) analysis was performed on exons of all protein-coding genes. For each gene, raw coverages were obtained by dividing the number of reads mapped to all (non-overlapping) exons by total exon length. Normalised gene coverages were then obtained by dividing raw gene coverages by the total number of reads passing mapping and quality filters from each sequenced library. In order to limit false positives in low coverage regions, genes showing a normalised coverage value < 1E-8 in at least one line were filtered out (12145 genes retained). As for RNA-seq, genes showing a CNV ≥ 1.5 fold (in either direction) in the four pairwise ‘resistant *vs* susceptible’comparisons were considered affected by CNV in association with resistance.

The divergence between the two copies of the P450 duplication was investigated through the detection of tri-allelic substitution sites in the IR0F resistant line. Substitution were called from reads mapped within a region encompassing the P450 duplication extended by 100 Kb on both sides (Chr1:271151705 - Chr1:271572082). Only substitutions showing a coverage > 40 reads and minimum allele frequency = 10 % were called, allowing the detection of 1401 and 1334 substitution sites within and outside the duplicated region respectively. Substitution sites were then considered as tri-allelic if they carried the reference allele together with two distinct variant alleles at the same position, each showing a frequency > 20 %.

### Long read sequencing and de novo assembly of the P450 duplication

The genomic architecture of the duplicated locus was further examined in the duplicated IR0F line versus the non-duplicated IR13 line. Both lines were submitted to long read sequencing using Oxford Nanopore technology. Genomic DNA was extracted from pools of 75 adult females using the Gentra Puregene Tissue kit (Qiagen) with 3 mL cell lysis buffer. Genomic DNA was quantified using Qubit DNA Broad Range assay (Qiagen) and quality-checked by agarose gel electrophoresis before adjusting concentration to 150 ng/µl. Long genomic DNA fragments were selected using the Short Read Eliminator kit (Circulomics). DNA libraries were prepared using 1 µg gDNA with the ligation sequencing kit SQK-LSK109 kit (Oxford Nanopore) following manufacturer’s instructions. Finally, 50 fmoles of the resulting DNA library were loaded on a R9.4.1 MinIon flow cell. Long read sequences were collected from 2 independent runs of 48 h for each line. Sequencing generated a total of 515K/1017K reads with an average length of 13.5/9.8 Kb, representing 3.4/4.8 fold genome coverage, for IR0F and IR13 lines respectively. Reads were mapped on the AaegL5 reference genome using Winnowmap (Jain et al., 2022) and alignments were visualised using IGV (Robinson et al., 2011). Finally, de novo genomes of the IR0F and IR13 lines were assembled using both short and long reads using MaSuRCA (Zimin et al., 2013). This de novo assembly resulted into 17346/ 13160 contigs, with a median size of 40/55 Kb for each line respectively. Transposable elements were identified using BLAST against the RepBase library (10/12/2021 version).

### Quantification of the P450 duplication

The detection of the P450 duplication was achieved through the quantification of *CYP6BB2* copy number by digital droplet PCR (ddPCR). Genomic DNA was extracted from pools of mosquitoes using the cetyltrimethylammonium bromide (CTAB) method (Collins et al., 1987), resuspended in nuclease-free water and quantified using Qubit DNA Broad Range assay (Qiagen). The duplex ddPCR quantification assay was based on the co-amplification of two target genes each detected with specific Taqman probes: the gene *CYP6BB2* (AAEL014893, Fam) was used to quantify the P450 duplication copy number while the gene *CYP4D39* (AAEL007808, Hex) always found as a single copy in various *Ae. aegypti* lines was used as control for normalisation of gDNA quantities (Faucon et al., 2015). Before amplification, gDNA samples were digested with XhoI for 15 min at 37°C directly within the ddPCR reaction mixture. This reaction mixture was partitioned into up to 20,000 nanoliter-sized droplets using the QX200 droplet generator (Bio-Rad) by mixing 70 μl of synthetic oil with 20 μl PCR mix. Then, 40 µL of the partitioned reaction was amplified for 40 cycles with an annealing temperature of 60 °C (see **Supplementary file 8** for protocol details and primers/probes). After amplification, the number of positive and negative droplets was quantified for both Fam and Hex channels using the QX200 droplet reader (Bio-Rad) and the positive/negative ratio was used to estimate the initial gDNA concentration of each gene assuming a Poisson distribution. After normalisation for gDNA quantity, *CYP6BB2* copy number were expressed as mean copy number (± 95

### Quantification of the Ile1011Met Kdr mutation frequency

The frequency of the Ile1011Met *Kdr* mutation was quantified on pools of mosquitoes using ddPCR. Genomic DNA was extracted as described above and digested with XhoI for 15 min at 37 °C before partitioning. The duplex TaqMan reaction mixture contained two amplification primers together with two Taqman probes labelled with Fam and Hex fluorophores corresponding to the Met and Iso alleles respectively (see **Supplementary file 8** for protocol and primer/probes details). The duplex reaction mixture was emulsioned as described above and amplified for 40 cycles with an annealing temperature of 59 °C. After droplet reading, the positive/negative ratio obtained for each channel was visualised as a scatterplot and was used to estimate the frequency of each allele assuming a Poisson distribution with 95 % CI.

## Acknowledgements

We thank the Institut Pasteur de La Guyane for providing the isofemale lines IR03 and IR13. We also thank Dr. Ademir Jesus Martins from the Fundación Oswaldo Cruz for a critical reading of the manuscript.

## Author contributions

JMB and JPD designed the study. TB, CH, JG, JC, MK, LN, TG, FL and JT performed experiments and analysed data. JF, FB, JV, JMB and JPD supervised data analysis. ID provided biological materials and data. JMB, JF and JPD acquired the funding. TB, JMB and JPD wrote and revised the paper.

## Data availability

All supplementary materials associated to this work have been deposited in Zenodo and are available at https://doi.org/10.5281/zenodo.14093547. 14093548. RNA-seq data have been deposited at EBI short read archive (SRA) under accession number E-MTAB-13325. DNA-seq short and long reads have been deposited at EBI short read archive (SRA) under accession number E-MTAB-13310.

## Ethical aspects

Mice were kept in the animal facility of the Biology department of the University of Grenoble-Alpes, approved by the French Ministry of Animal Welfare (agreement no. B 38 421 10 001) and used in accordance with the laws of the European Union (directive 2010/63/EU). The use of animals for this study was approved by the ComEth Grenoble-C2EA-12 ethics committee mandated by the French Ministry of Higher Education and Research (MENESR). The study was conducted in accordance with the ARRIVE guidelines.

## Supplementary materials

**Supplementary file 1 (.tiff image). Bioassays data from F1 and F2 offspring used to investigate the mode of inheritance of the resistance phenotype**. The reciprocal controlled crosses used to generate F1 and F2 individuals used for bioassays are indicated. Deltamethrin exposure conditions were 0.03 % deltamethrin for 30 min for females and 0.015 % deltamethrin for 25 min for males in order to account for their inherent lower insecticide tolerance. Mortality rates are indicated as mean ± SD. Distinct letters indicate significantly distinct means (Kruskal-Wallis test followed by post hoc pairwise Wilcoxon test with Bonferroni-Holm correction, N = 5, p ≤ 0.05).

**Supplementary file 2 (.xlsx file). RNA-seq and CNV data sets**. All protein coding genes showing a significant differential transcription (sheet ‘RNA-seq’) or a significant differential exon coverage (sheet ‘CNV’) are shown. See methods for filtering conditions and thresholds. Gene descriptions beginning with a # were manually inferred from Blast X result.

**Supplementary File 3 (.tiff image). Genomic architecture of the *Kdr* locus in the IR0F resistant line**. On chromosome 3, the 3’ part of the VGSC gene, including the 21 last exons, is duplicated. Black bars indicate the position of the coding exons. The duplication is 125 Kb long, and is imperfect: the two copies, R and R’ mostly differing in their intronic sequences. The duplication breakpoints, left (LB) and right (RB) are located at 315,905,811 bp and 316,030,250 bp on the reference genome AaegL5. The 1011Met codon is located on the upstream copy (R) and followed by a B type intron. The 1011Ile codon is on the downstream copy and followed by an A type intron (Martins et al., 2013). RNA-seq data supported the sole expression of the 1011Met allele.

**Supplementary file 4 (.xlsx file). Long reads overlapping the P450 duplication breakpoints**.

**Supplementary file 5 (.docx file). Data supporting the identification of the transposable elements flanking the P450 duplication**.

**Supplementary file 6 (.docx file). RNA interference experiment**. This document includes detailed results and methods.

**Supplementary file 7 (.tiff image). Proposed evolutionary steps leading to the observed duplication architecture**. Theoretical structures and those inferred from short and long reads data are indicated. The hAt-like and PiggyBac-like transposons are shown in purple and green respectively. Transposition events are shown as dashed lines. The second hAt-like transposition event would have created an incomplete copy of the hAT transposon, either directly or through a posterior deletion. Recombination events are shown as blue crossed lines. Stars indicates junctions that were confirmed by short reads data.

**Supplementary File 8 (.docx file). Digital droplet PCR assays used to quantify the P450 duplication and the frequency of the Ile1011Met *Kdr* mutation**. This document provides all technical details related these ddPCR assays including amplification primers, TaqMan probes, PCR conditions and data interpretation.

