## Supplementary figures and images for "A genomic duplication spanning multiple P450s contributes to insecticide resistance in the dengue mosquito *Aedes aegypti*"

### Supplementary file 1

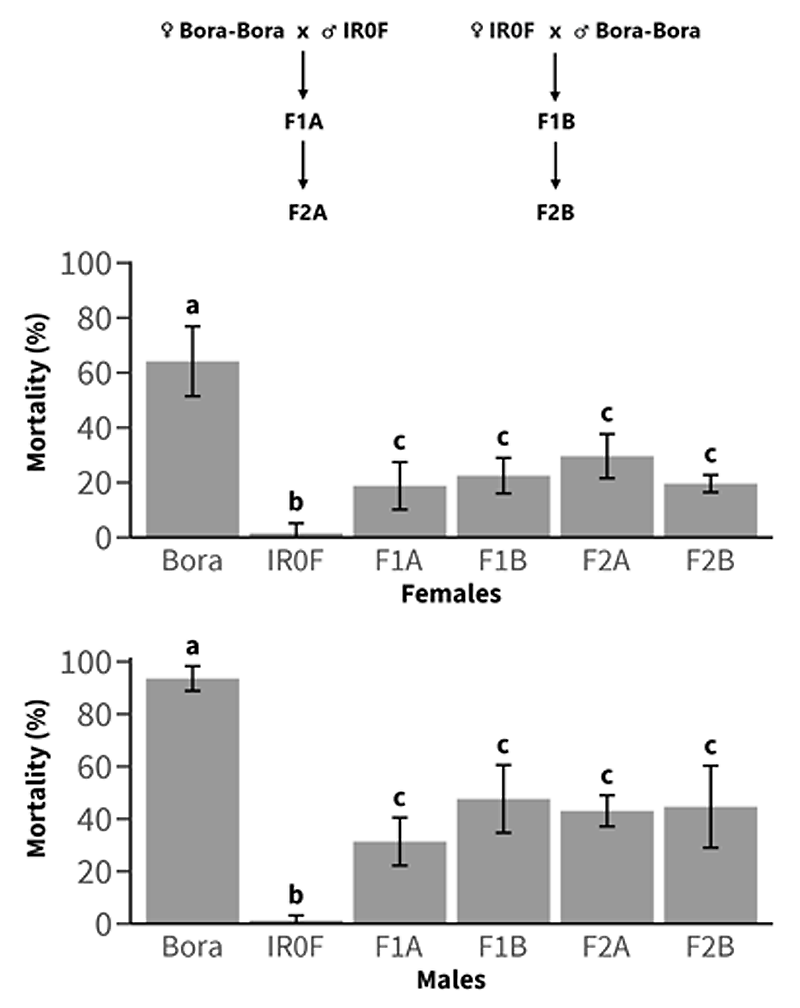

### Supplementary file 3

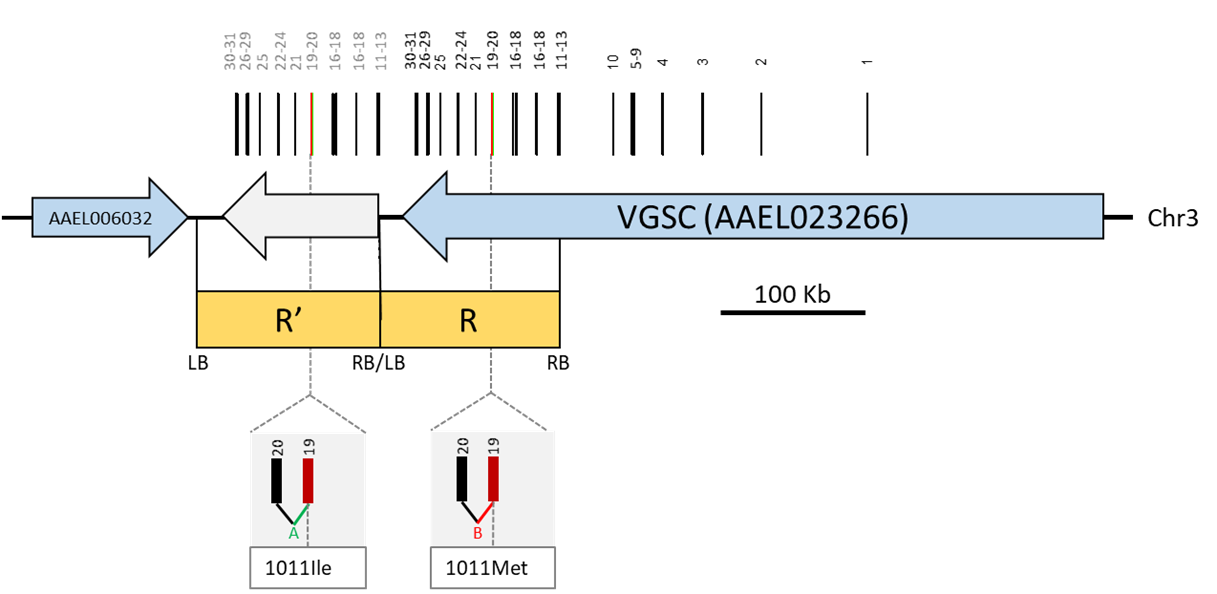

### Supplementary file 7

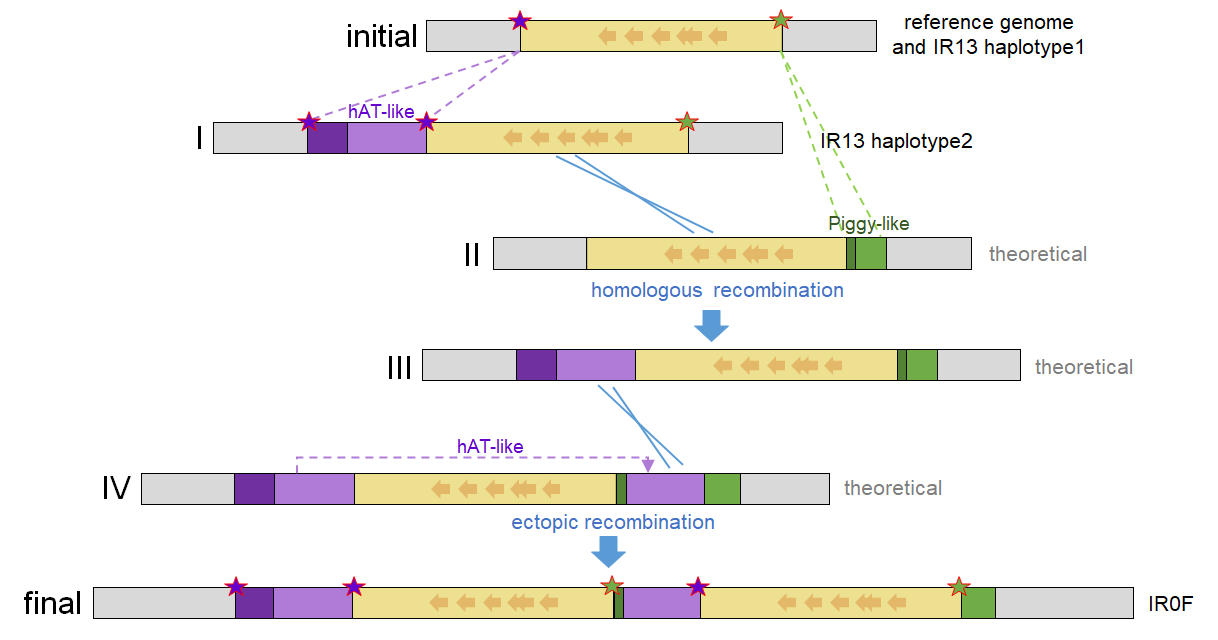
