## Supplementary file 5 for "A genomic duplication spanning multiple P450s contributes to insecticide resistance in the dengue mosquito *Aedes aegypti*"

### Supplementary File 5. Description of the transposons flanking the P450 duplication

#### I- PYL: a PiggyBac-like transposon at the right breakpoint of the duplication

On its right side, the CYP6 cluster duplication was flanked by a 2.4 Kb insertion, hereafter named PYL (for PiggyBac-like). PYL sequence ends with 23 bp terminal inverted repeats (TIR) in agreement with a DNA transposon, and two >99% related sequences were found inserted elsewhere in the reference AaegL5 genome (Figure 1). The PYL Terminal Inverted Repeats (TIRs) were however flanked by a target site duplication (TSD) of only 2 bp (Figure 1), while most DNA transposons show longer TSDs (Schulman 2013). A feature reminiscent of the PiggyBac transposon is the presence close to the transposon ends of several internal direct and inverted repeats (Chen et al. 2020; Figure 1).

```
Empty site    AACCACAGAT..................................................
PYL           AACCACAGATacagtgggattccgtttttggcatgctccgtttttggcatgctccgattt----
Chr1_236Mb    TGAAATAGATacagtgggattccgtttttggcatgctccgtttttggcatgctccgattt----
Chr2_33Mb     CTTTATTTATacagtgggattccgtttttggcatgctccgtttttggcatgctccgattt----
                        >>>>>>>>>>>>>>>>>>>>>>>
                         .        .         .         .         .         .
                         1        10        20         30        40        50


Empty site                             ....................................................................ATATGGACTG
PYL                                    ----acggttccgtttttggcaactgaaaatttcaactttgtttgccaaaaacggaatcccactgtATATGGACTG
Chr1_236Mb                             ----acggttccgtttttggcaactgaaaatttcaactttgtttgccaaaaacggaatcccactgtATCAACAAAA
Chr2_33Mb                              ----acggttccgtttttggcaactgaaaatttcaactttgtttgccaaaaacggaatcccactgtATTGGACTTT
                                                                             <<<<<<<<<<<<<<<<<<<<<<<
                                           .         .         .         .         .         .         .
                                          2355      2365      2375      2385      2395      2405      2414
```

***Figure 1. The PYL terminal sequences.*** *The transposon sequence is displayed in lowercase, and the surrounding sequence in uppercase. The ATAT sequence present at empty sites is shown in red. Blue letters denote the 23 bp TIRs. A perfect 24 bp direct repeat is present near both transposon ends and shown in green. Underlined is a smaller perfect repeat. From top to bottom: the empty site sequence found both in AaegL5 and in IR13, the PYL element found in the IR0F line, and two sequences >99% homologous from AaegL5 (at Chr1:236484856-236484966 and Chr2:33998279-33998389, respectively).*

In search for transposon activity, we isolated sequences flanked by the 23 bp PYL TIR sequences in opposite orientations, and identified 996, 933 and 1089 matches in the genomic assemblies of IR13, IR0F and AaegL5, respectively, with a median size of 2.7 Kb. For the IR13 line, 193 of these elements were mapped unambiguously on the reference genome (homology extending over > 1Kb on both sides in a non-repeated region). Of these, 29 elements corresponded to perfect empty site positions with no transposon in the AaegL5 genome, supporting recent transposition events in the PYL transposon family by a cut-and-paste mechanism.

The PiggyBac transposon, originally isolated from the *Trichoplusia ni* butterfly, targets the TTAA sequence, which is duplicated upon insertion and restored as single copy upon excision (Cary *et al.* 1989; Chen *et al.* 2020). By contrast, PYL related elements displayed a shorter target specificity, namely T/A (Figure 2), implying a distinct transposition mechanism.

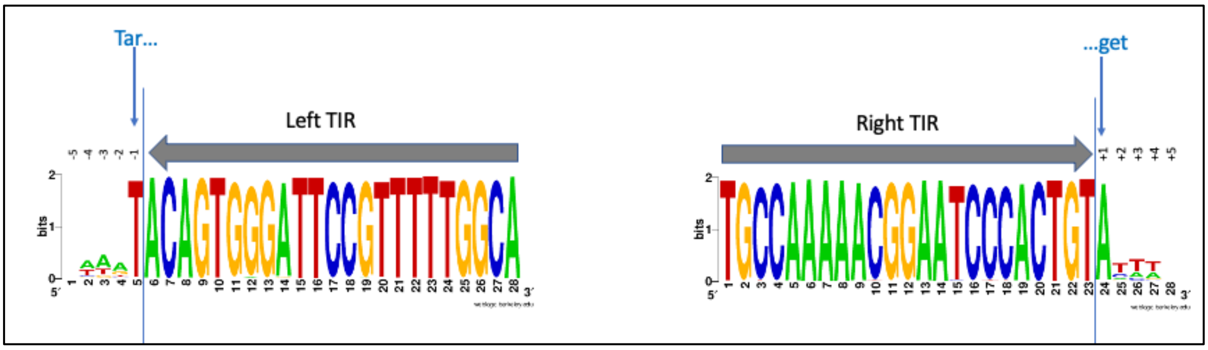


***Figure 2. Sequence conservation of the left and right TIRs with the 5 flanking bases of target sequence.*** *The 23 bp PYL TIR sequence was used as query in a Blast of the genomic assembly, and sequences sandwiched between a left TIR on the left and a right TIR on the right were then extracted as PYL related. Consensus sequences are derived from the 982 PYL-related sequences found in IR13: note the absence of a conserved duplicated sequence flanking the transposon TIRs. Consensus were produced at weblogo.berkeley.edu.*

PYL contains no open reading frame and therefore defines as a non-autonomous transposon. However, one PYL-related element in the IR13 assembly (scaffold 7180000050065:279965-295286) was identified with a predicted protein homology to the *Trichoplusia ni* PiggyBac transposase and named PYLp (Figure 3). Homologies were found in the two PYLp ORFs, suggesting they are spliced together before translation.

In light of these data, the PYL transposon likely belongs to a family of class II transposons actively moving through the *Ae. aegypti* genome and related to Piggy-Bac, but likely transposing through a somewhat different mechanism.

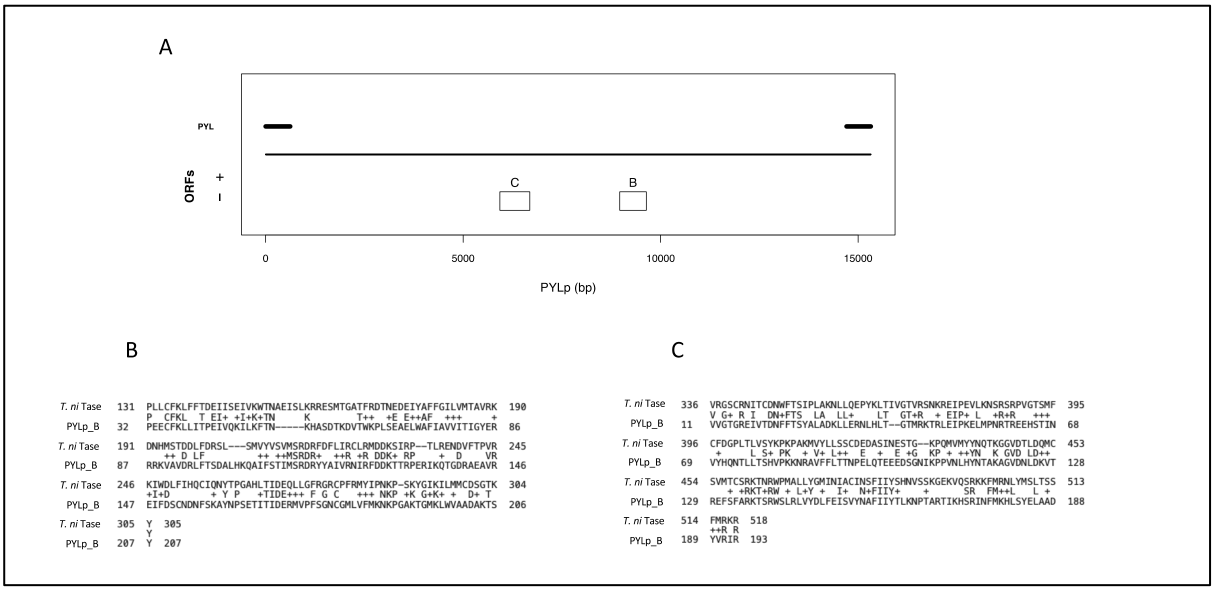


***Figure 3.*** *PYLp****,*** *a PYL related element showing homology to the piggyBac transposase.* ***A****: Motives in the PYLp element. Thick lines indicate DNA homology to PYL, which extends beyond TIRs. White boxes indicate the two ORFs, located on the minus strand.* ***B and C****, protein homologies of the translated orfB and orfC to the T. ni transposase (pdb 6X68).*

### II- A hAT transposon at the left breakpoint of the duplication

On the left side, the duplication was flanked by a 9040 bp insertion, hereafter named CYP6-hAT, ending with 15 bp terminal inverted repeats (TIR), and flanked by an 8 bp target site duplication (Figure SX4 A). Close to the TIRs, four subterminal instances of the *TAATCgatta* palindrome were found. Both the length of the TSD and the presence of subterminal repeats suggest that this element belong to the hAT family of cut-and-paste DNA transposons.

Three nearly identical elements (>95% identity on the whole length) were found in the IR0F genome assembly, and two in the reference genome (Figure 4B). In addition, a genomic search for sequences flanked by the same TIRs allowed to define a family containing 157, 143 and 149 elements in the IR13, IR0F and the reference AaegL5 genome assemblies respectively, with a median size of 8,3 Kb. A perfect 8 bp repeat (*i.e.* a target site duplication) is flanking 78% of these elements, and they contain on average more than seven internal instances of the *TAATCgatta* palindromic sequence, all within 200 pb of either TIR. Among the 37 insertion sites identified in the IR13 line and mapped unambiguously on the reference genome, 9 correspond to empty sites in the AaegL5 genome, supporting recent transposition events.

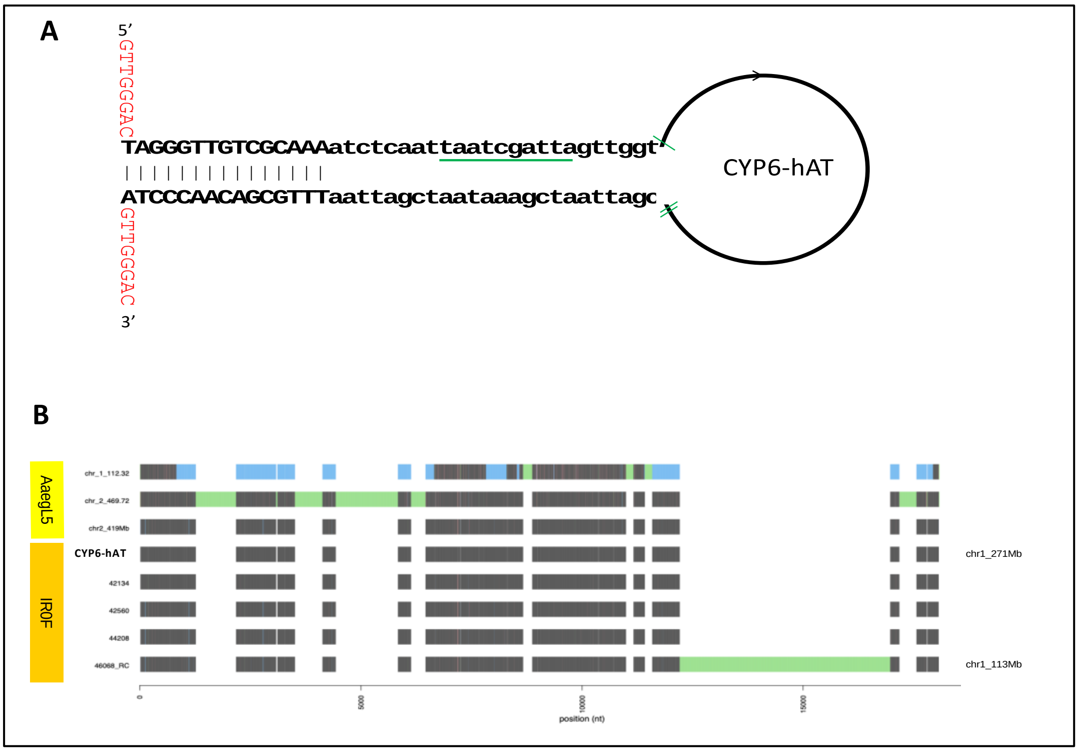


***Figure 4:** [**A**] **Structure of the CYP6-hAT element**. The 8 bp target site duplication (TSD) is shown in red, and the 3' repeat marks the left breakpoint of the CYP6 cluster duplication in IR0F. The first instance of the TAATCgatta palindrome is green underlined, the green ticks indicate other occurrences. [**B**] **DNA elements most closely related to CYP6_hAT**. Black boxes represent perfect identity to the CYP6_hAT sequence, blue and green boxes represent deletions and insertions respectively. The three first elements are in the AaegL5 reference genome (positions indicated). The five subsequent elements were found in the IR0F assembly, and their position is given on the right when non-ambiguous (i.e. flanking sequences from both sides contiguous in the reference genome).*

CYP6-hAT has no coding capacity (*i.e.* is a non-autonomous transposon). We searched for *Ae. aegypti* hAT family members with at least one internal open reading frame (ORF) longer than 500 bp, and identified 33, 29, and 39 such elements in IR13, IR0F and AaegL5, respectively. Among those, we identified one possibly autonomous element in IR13, hereafter hATp (Figure 5A) : hATp shows protein homology to Hermes, a hAT transposase from *Musca domestica* (Hickman et al. 2014) (Figure 5B). This homology spans two ORFs, m5 and m4, that are likely exons of a same gene and covers residues involved in Hermes TIR DNA binding (Hickman et al. 2014). In addition, a BED_Zn finger motif is present at the N terminus of the m5 ORF, alike in hAT transposases from other species where this domain is also known to contribute to TIR DNA binding (Jiang et al. 2016). Interestingly, the hATp element shows a Russian Doll architecture, such that this transposon could disseminate both using its own TIRs and a transposase as a cargo gene of a Gypsy retrotransposon (Figure 5A).

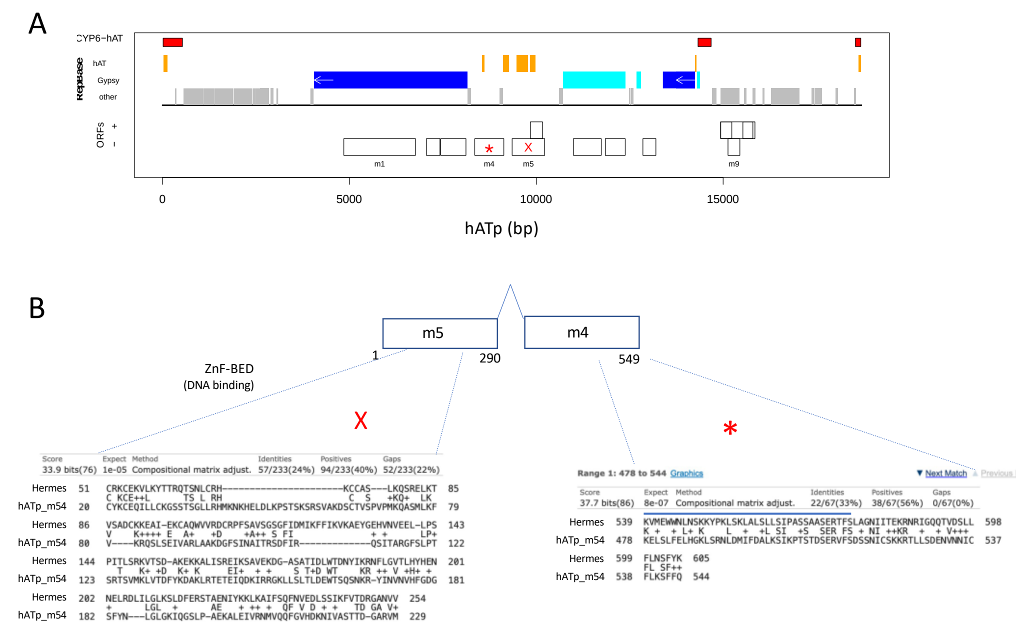


**Figure 5. hATp, a putative autonomous hAT element related to CYP6-hAT found in the IR13 genomic assembly.** *This hATp was identified from scaffold 7180000034851 at positions 154560-173223 of our de novo assembly.* ***[A], DNA architecture of hATp****. Red boxes indicate areas conserved with CYP6-hAT (note that homology extends beyond the TIRs), found by BLAST. Orange, blue and grey boxes indicate homology blocks found by Repbase to hAT transposons, Gypsy retrotransposons, and other transposable elements, respectively. The Gypsy-232 related sequences appear in dark blue, with arrows indicating the LTRs. White boxes indicate open reading frames. Red symbols x and * indicate homology to Hermes transposase.* ***[B], Protein homologies*** *to Hermes hAT-like transposase from Musca domestica (Q25438, 612 aa)****.*** *The ORFs m5 and m4 from hATp were assumed to be spliced together and translated into a putative m5-4 protein; amino acid numbering assumes splicing with no coding codon loss. Residues in purple boxes are known to interact with the Hermes terminal DNA sequence. The predicted N-terminal BED_Zn finger motif is shown in yellow. The blue line indicates the extent of domain III, used for multiple alignment in Figure 6.*

The CYP6-hAT TIR sequence shows some base conservation with the TIRs of the hAT superfamily (Figure 6A). Also, alignment of transposase protein sequences in a conserved C-terminal area (domain III, Rubin et al. 2001) indicates strong differences of the *Ae. aegypti* hATp gene product to the known insect hAT transposases: out of 39 residues, Hermes, Hobo, Hermit and Homer share 16 identical positions with each other, whereas hATp shares only 4 positions with them (Figure 6B). In light of these data, the CYP6-hAT transposon appears to be an actively moving member of the hAT in *Ae. Aegypti* genome.

A

| | | | | |
|---|---|---|---|---|
| Ac | TAGGGATGAAA | Zea mays | Plants | 168402 |
| Tam3 | TAAAGATGTGAA | A. majus | Plants | 16063 |
| GI4314354 | TAGGGGTGTCAAAA | A. thaliana | Plants | 4314354 |
| CYP6-hAT/hATp | TAGGGTTGTCGCAAA | Ae. aegypti | Insects | |
| Hermit | CAGAGATGTGCATG | L. cuprina | Insects | 726316 |
| Hobo | CAGAGAACTGCA | D. melanogaster | Insects | 157606 |
| Hermes | CAGAGAAC | M. domestica | Insects | 514387 |
| Homer | CAGAGATCTGCA | B. tryoni | Insects | 4106909 |
| Restless | CAGAGTGCGTAATC | T. inflatum | Fungi | 1542944 |

B

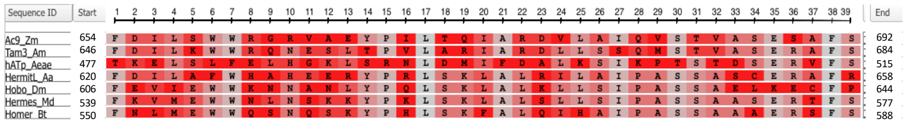

***Figure 6. Multiple alignments of hAT sequences. [A],*** *Alignment of the terminal inverted repeats of plant, insect and fungal hAT transposons: GI accessions are indicated. Bold letters indicate conserved bases between CYP6-hAT TIR and other hAT TIRs, retrieved from Rubin et al. 2001).* ***[B], Alignment of hAT transposases*** *in a conserved C-terminal region (domain III, Rubin et al 2001). Note that hATp-m54 is only distantly related to other insect transposases. Ac9 ,TRAC9 (Zea mays); Tam3, Q38743 (Antirhynum majus); Hermes, Q25442 (Musca domestica); Hobo, P12258 (Drosophila melanogaster); Hermit, KAH7703716.1 (Aphelenchus avenae); Homer, O96691 (Bactrocera tryoni). Alignment viewed on https://www.ncbi.nlm.nih.gov/projects/msaviewer using frequency based preferences: conserved residues appear in grey.*
