## Supplementary file 6 for "A genomic duplication spanning multiple P450s contributes to insecticide resistance in the dengue mosquito *Aedes aegypti*"

**Supplementary file 6: RNAi-silencing of CYP6 genes carried by the duplication**

Results:

RNA interference through intrathoracic dsRNA injection was used to investigate the relative importance of the different CYP6 genes carried by the duplication in the deltamethrin resistance phenotype. Only the first five CYP6 of the duplicated cluster were targeted as CYP6CC1 showed a very low expression level in adult females as measured from RNA-seq.

Investigating silencing efficiency and specificity using RT-qPCR showed that dsRNA injections lead to an incomplete silencing of the targeted CYP6s with a silencing efficiency ranging from 42.2% for AAEL017061 to 69.7% for AAEL026582 (**Figure 1**). A good overall silencing specificity was observed for most targeted CYP6s despite their high sequence homology. However, a significant cross-silencing effect was observed for the dsRNA targeting AAEL014891v1 (CYP6P12v1) toward AAEL014891v2 (CYP6P12v2) and in a lesser extent for the dsRNA targeting AAEL026852 toward AAEL017061.

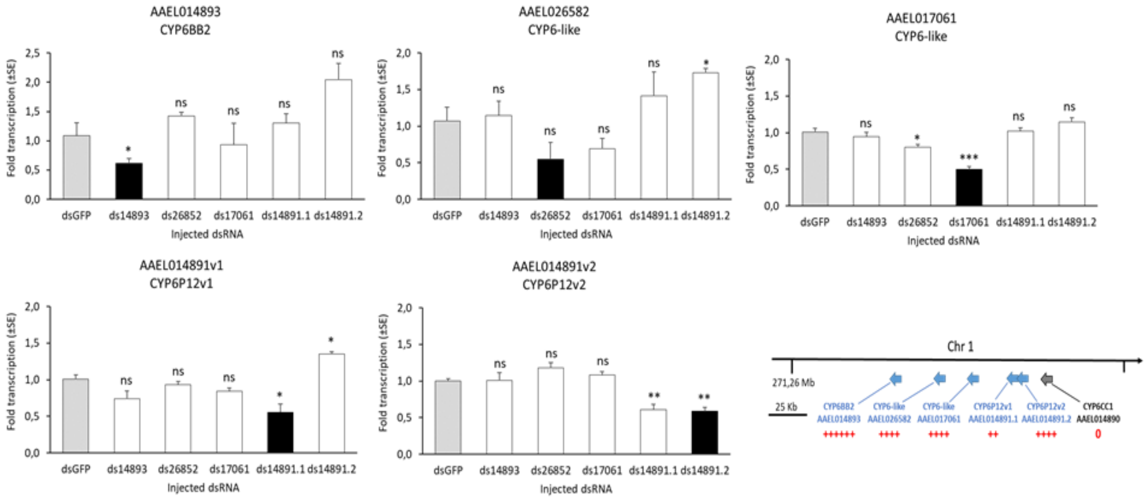


**Figure 1:** Knock down efficiency and specificity of dsRNA constructs. Silencing efficiency was assessed using RT-qPCR through the comparison of transcription levels between mosquitoes injected with dsGFP (grey) and those injected with each dsCYP6 construct (black). Silencing specificity was assessed by comparing the transcription level of each target CYP6 gene (black) between mosquitoes injected with its respective dsCYP6 construct and those injected with other dsRNA constructs (white). DsRNA construct names reflect the accession number of their respective target gene. Comparisons were made using Student's t-tests (N=4, * $P<0.05$, ** $P<0.01$). The right panel shows the position of each gene within the CYP6 cluster together with its basal transcription level in adult females (red). The gene CYP6CC1 at the end of the duplicated CYP6 cluster was not targeted due to its very low expression level.

Comparative deltamethrin bioassays between mosquitoes injected with dsGFP (controls) and dsRNA constructs targeting each CYP6 gene revealed a limited effect of CYP6 silencing on deltamethrin survival (**Figure 2**). The mortality observed in mosquitoes injected with the dsGFP construct varied from 17 to 32 % across 5 injection experiments. Overall, the mean mortality rates observed upon injection of a dsRNAs targeting each CYP6 gene were not significantly different from the controls. However, a significant higher mortality was observed for mosquitoes injected with dsRNA constructs targeting CYP6BB2 and CYP6-like AAEL023582 as compared to mosquitoes injected with dsRNA constructs targeting the three other CYP6 genes (AAEL017061, CYP6P12v1 and CYP6P12v2).

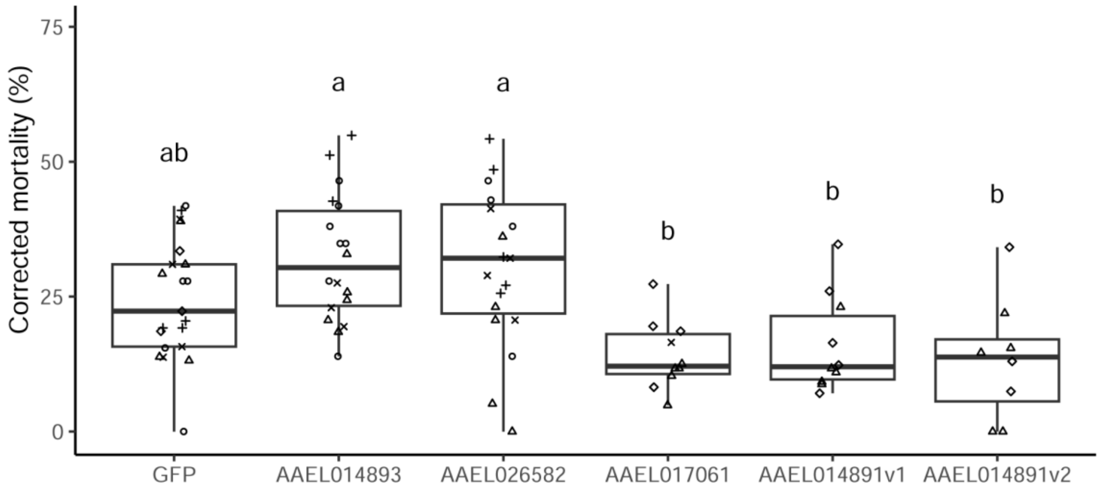


**Figure 2.** Comparative deltamethrin mortalities of IR0F females injected with dsRNA targeting the exogenous gene GFP (controls) or CYP6 genes carried by the duplication. Data were obtained from five injection experiments using different mosquito batches. Mortality rates obtained from the different injection experiments were normalized according to the mean mortality obtained for dsGFP controls. Each point represents a WHO tube (mean= 16.5 injected mosquitoes), and different shapes represent different injection experiments. The mean mortalities were compared with a Kruskal-Wallis rank sum test ($p<10^{-4}$), followed by pairwise Wilcoxon tests with Holm correction of p-values; a and b represent significantly distinct means ($p<0.05$).

Methods

The primers for dsRNA constructions were designed on NCBI Primer Blast to amplify specific products for each gene of interest with a length of 500-600 bp and T7 promoter sequence (5' TAATACGACTCACTATAGGG 3') at the 5' end of both forward and reverse oligos (**Table 1**). PCR amplification was carried out using cDNA from the IR0F line, using Phusion High-Fidelity DNA Polymerase (New England Biolabs) in 50 µl reactions following manufacturer's instructions in the following cycle: 30 seconds at 98°C for initial denaturation, 35 cycles of 10 seconds at 98°C for denaturation, 30 seconds at 62-64°C for annealing and 15 seconds at 72°C for extension and a final hold at 72°C for 10 minutes. Amplified products were subjected to PCR purification using Nucleospin™ Clean-up Kit (Macherey-Nagel). Synthesis of dsRNAs was carried out using HiScribe™ T7 High Yield RNA Synthesis Kit (New England Biolabs) and the purification was performed using MEGAclear™ Transcription Clean-Up Kit (Ambion). Elution was performed with Nuclease-free water and the concentrations were estimated using nanodrop ND1000 (Thermofisher). All dsRNAs were then diluted to 3 µg/µl for injections.

Two days-old non-blood fed females of the IR0F were anaesthetized with $C0_2$ and subjected to nano-injections using Nanoject II nano-injector (Drummond Scientific Company). Up to 70 nl of dsRNA was injected intrathoracically, underneath the wing resulting in approximately 200 ng injected dsRNA per mosquito. A 500 bp dsRNA prepared from *Green fluorescent protein* (GFP) non-endogenous gene (mentioned as dsGFP) was used as control. Injected mosquitoes were then kept in small cages with cottons impregnated with a 1% honey solution as food source.

Seventy-two hours post injection, four replicates of 5-8 mosquitoes for each dsRNA were used to assess the knock down efficiency and specificity of each dsRNA using RTqPCR. Total RNA extractions were performed using TRIzol™ following manufacturer's instructions and total

RNA was eluted in nuclease-free water and quantified using a nanodrop ND1000. A DNase I treatment (Invitrogen) was carried out to remove potential genomic DNA contamination. Two µg of DNase-treated RNA were used for reverse transcription using oligodT and Superscript$^{TM}$ III First Strand Synthesis System (Invitrogen) following manufacturer's instructions. Specific RT-qPCR primers were designed out of the dsRNA target regions for each gene of interest using NCBI Primer Blast. Primers were designed, with 60°C melting temperatures, amplicons of 80-180bp and where possible spanning exon junctions. Quantitative PCR reactions were performed on a CFX-96 real time system (BioRad) with each 25 µl reaction containing: 12.5 µl iQ SYBR Green Supermix, 0.3 µM of each primer and 5 µl of 1:25 diluted cDNA. A standard curve was used to estimate PCR efficiency using a mix from all the cDNAs diluted from 1:5 to 1:625 (90%-120% for all primer pairs). All qPCR reactions were performed in duplicates with the following cycles: 3 min at 95°C, 40 cycles of 15 seconds at 95°C and 30 seconds at 60°C. Relative transcription levels were obtained for each gene using the two control genes RPS7 and RPL8. Silencing efficiency (RNAi knock-down) for each dsRNA was estimated by comparing the relative transcription level of each target gene between dsRNA-injected and dsGFP-injected mosquitoes. Since the P450s carried by the duplication belong to the same CYP6 subfamily, off-target effects were assessed by comparing the relative transcription level of the CYP6 gene targeted by dsRNA with the four other CYP6 genes. Pairwise statistical comparisons of transcription levels were performed using Student's t-test (N=4).

**Table 1.** Primers used for the synthesis of RNAi constructs and RT-qPCR

| Target gene | Expression level (females) | dsRNA construct primers | | Silencing efficiency (%) | Efficiency P value | RT-qPCR primers | RT-qPCR efficiency (%) |
|---|---|---|---|---|---|---|---|
| AAEL014893 (CYP6BB2) | high | 14893_ds2 | **(T7)**CGGTTTGACGGGGTAAGGTT<br>**(T7)**CCTGCAGGGATGGTGAAGTT | 64,5 | 0,0034 | AAGGTTGCAGGATGTCCAGG<br>TCATTCGACAACGCATCCCA | 96.5 |
| AAEL26582 (CYP6-Like) | moderate | 26853_ds3 | **(T7)**CTGAACGAGTTGGCTGCTCA<br>**(T7)**CTTCCACCTCGGCAAACTGT | 69,7 | 0,0013 | AAACCTGGACGTTGGGAGAC<br>CATTGGTCTTCTCGCGGTGA | 100 |
| AAEL017061 (CYP6-like) | moderate | 17061_ds2 | **(T7)**CGTACCATACGTTCCCGGTT<br>**(T7)**GACTCTGTCGGCAATGTCCA | 42,2 | 0,0002 | TTGCTGAGTCGGTTACGCTT<br>CACATCCAACCAAAGTCCGC | 100 |
| AAEL014891v2 (CYP6P12v2) | moderate | 14891.2_ds3 | **(T7)**GGTGTTCGAGCAGAGGATGT<br>**(T7)**GGGAACCGTGTAGTCCGATG | 64,8 | 0,0006 | TGTGAATGAAACCCTTCGCAAG<br>GCAACGCATAGATCGGTATGG | 100 |
| AAEL014891v1 (CYP6P12v1) | low | 14891.1_ds2 | **(T7)**CCTGGACAAGAAGCGGAGTT<br>**(T7)**AGCATCTGCCCTACGGATTG | 47 | 0,0332 | GCTGAAGCTCAAAGCCACTC<br>GTTGTTCCGTTCTCGGTGGT | 98,6 |

The impact of the silencing of each CYP6 gene on deltamethrin resistance was then examined by injecting females of the IR0F line with each CYP6 dsRNA construct or with the dsGFP construct as control. Considering the high number of injected mosquitoes and the time required for injections, five distinct experiments were performed, each targeting from two to five distinct CYP6 genes in addition to the control. For each experiment, approximately 100 two days-old females were injected with each dsRNA and allowed to recover for 72h as described above before being used for comparative deltamethrin bioassays. Batches of 10-20 injected mosquitoes were exposed to 0.03% deltamethrin for 90 min using WHO test tubes as previously described and mortality was recorded 24h post exposure. In order to standardize bioassay data across distinct experiments, % mortalities obtained during each experiment were normalized according to the mean mortality obtained for dsGFP controls.
