## Supplementary file 8 for "A genomic duplication spanning multiple P450s contributes to insecticide resistance in the dengue mosquito *Aedes aegypti*"

### Supplementary file 8: digital droplet PCR assays

This document provides all technical information related to the digital droplet PCR TaqMan assays used to quantify CYP6BB2 copy number and to quantify the frequency of the Ile1011Met *kdr* mutation.

#### 1. Quantification of *CYP6BB2* copy number using ddPCR

##### 1.1. Primers and probes

| Gene | Accession number | Primer name | Primer sequence (5' to 3') |
|---|---|---|---|
| ***CYP4D39* (control)** | AAEL007808 | F_CYP4D39 | AGTCCTGGAAGTTCTGCACG |
| | | R_CYP4D39 | AAGGCGACTTTCCGACGAAT |
| | | CYP4D39(Hex) | **[HEX]**AAGGAGGCAAACCCCGATAA**[BHQ1]** |
| ***CYP6BB2*** | AAEL014893 | F_CYP6BB2 | AGTTCAAGGGCCGAGGATTG |
| | | R_CYP6BB2 | CGGATCCACGAAAATTCCGC |
| | | CYP6BB2(Fam) | **[6FAM]**GCTGGTTATCGATCCGGAGA**[BHQ1]** |

##### 1.2. ddPCR reaction Mix

| Reagent | Initial concentration | Volume (µl) |
|---|---|---|
| ddPCR Supermix for Probes (No dUTP) | 2X | 12.5 |
| R_CYP6BB2 | 10 µM | 1.125 |
| F_CYP6BB2 | 10 µM | 1.125 |
| CYP6BB2* | 10 µM | 0.625 |
| R_CYP4D39 | 10 µM | 1.125 |
| F_CYP4D39 | 10 µM | 1.125 |
| CYP4D39* | 10 µM | 0.625 |
| RNAse-free water | - | 2.39 |
| Restriction enzyme XhoI | 10 U/µL | 0.25 |
| rCutSmart buffer | 10 X | 0.11 |
| gDNA | 5 ng/µL | 4 |
| | **TOTAL VOLUME** | **25** |

1.3. ddPCR conditions

| Step | Temperature | Duration |
|---|---|---|
| 1. Polymerase activation | 95 °C | 10 minutes |
| 2. Denaturation | 95 °C | 10 seconds |
| 3. Annealing + Extension | 60 °C | 45 seconds |
| 4. Repeat (2. + 3.) X 39 | | |
| 5. End | 4 °C | 5 minutes |
| 6. Polymerase inactivation | 90 °C | 5 minutes |
| 7. Post-PCR waiting time | 4 °C | ∞ |

#### 2. Quantification of *kdr* Ile1011Met allele frequency using ddPCR

##### 2.1. Primers and probes

| Gene | Allele | Primer name | Primer sequence (5' to 3') |
|---|---|---|---|
| Voltage-Gated Sodium Chanel Para (AAEL023266) | | F_I1011M | TCCAATTTACTTTCAGTCAGC |
| | | R_I1011M | TGCTTGTGGGTGACG |
| | 1011_Iso (wild type) | 1011Iso(Hex) | **[HEX]**CTAGATTTCCTATCACTACGGTGG**[BHQ1]** |
| | 1011_Met (mutated) | 1011Met(Fam) | **[6FAM]**ACTAGATTTCCCATCACTACGGT**[BHQ1]** |

##### 2.2. ddPCR Mix

| | Initial concentration | Volume (µl) |
|---|---|---|
| ddPCR Supermix for Probes (No dUTP) | 2X | 12.5 |
| R_I1011M | 10 µM | 1 |
| F_I1011M | 10 µM | 1 |
| 1011Iso(Hex) | 10 µM | 0.25 |
| 1011Met(Fam) | 10 µM | 0.25 |
| RNAse-free water | - | 5.64 |
| Restriction enzyme XhoI | 10 U/µL | 0.25 |
| rCutSmart buffer | 10 X | 0.11 |
| gDNA | 5 ng/µL | 4 |
| | **TOTAL VOLUME** | **25** |

#### 2.3. ddPCR conditions

| Step | Temperature | Duration |
|---|---|---|
| 1. Polymerase activation | 95 °C | 10 minutes |
| 2. Denaturation | 95 °C | 10 seconds |
| 3. Annealing + Extension | 59 °C | 45 seconds |
| 4. Repeat (2. + 3) X 39 | | |
| 5. End | 4 °C | 5 minutes |
| 6. Polymerase inactivation | 90 °C | 5 minutes |
| 7. Post-PCR waiting time | 4 °C | ∞ |

#### 2.4. Scatterplots profiles obtained from the observed Iso1011Met genotypes

Y axis = Fam channel (Met allele), X axis = Hex chanel (Ile allele)
The resistant allele is duplicated (Ile-Met). The Wild type allele is in single copy (Ile)
Grey: Ile-/Met- droplets, Blue: Ile-/Met+ droplets, Green: Ile+/Met- droplets, Orange: Ile+/Met+ droplets

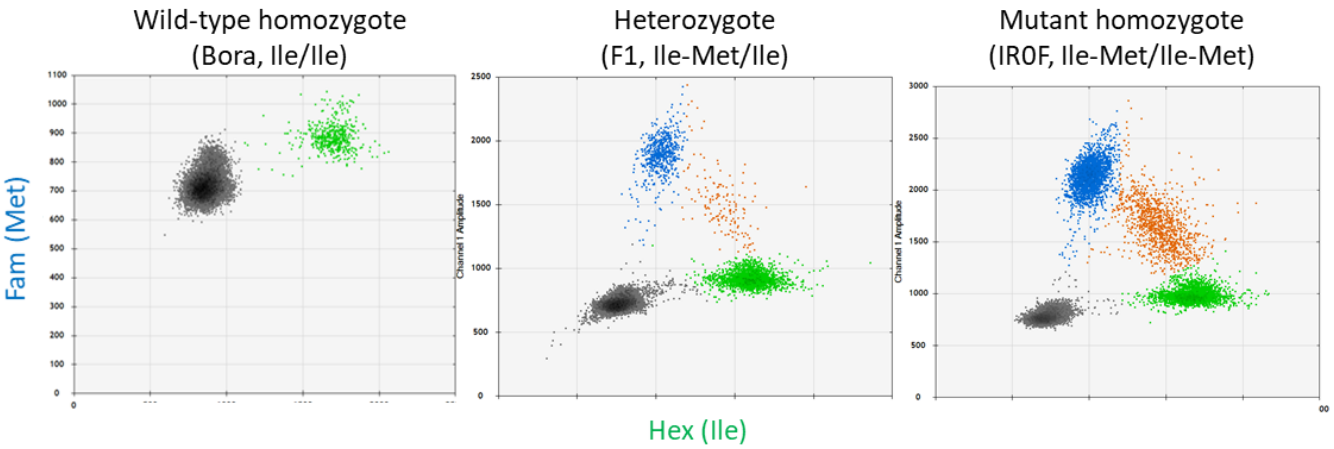
